# The mobility of interfaces between monomers in dimeric bovine ATP synthase participates in the ultrastructure of inner mitochondrial membranes

**DOI:** 10.1101/2020.09.18.303636

**Authors:** Tobias E. Spikes, Martin G. Montgomery, John E. Walker

## Abstract

The ATP synthase complexes in mitochondria make the ATP required to sustain life by a rotary mechanism. Their membrane domains are embedded in the inner membranes of the organelle and they dimerize via interactions between their membrane domains. The dimers form extensive chains along the tips of the cristae with the two rows of monomeric catalytic domains extending into the mitochondrial matrix at an angle to each other. When the interaction between membrane domains is disrupted in living cells, the morphology of the cristae is affected severely. By analysis of particles of purified dimeric bovine ATP synthase by cryo-electron microscopy, we have shown that the angle between the central rotatory axes of the monomeric complexes varies between *ca*. 76^°^ and *ca*. 95^°^. Some variations in this angle arise directly from the catalytic mechanism of the enzyme, and others are independent of catalysis. The monomer-monomer interaction is mediated mainly by j-subunits attached to the surface of wedge shaped protein-lipid structures in the membrane domain of the complex, and the angular variation arises from rotational and translational changes in this interaction, and combinations of both. The structures also suggest how the dimeric ATP synthases might be interacting with each other to form the characteristic rows along the tips of the cristae via other inter-wedge contacts, moulding themselves to the range of oligomeric arrangements observed by tomography of mitochondrial membranes, and at the same time allowing the ATP synthase to operate under the range of physiological conditions that influence the structure of the cristae.

## INTRODUCTION

The ATP synthase complexes embedded in the inner membranes of mitochondria are rotary machines that make the ATP required to sustain life (Walker, 1998). They generate rotational torque directly from the trans-membrane proton motive force (pmf) generated by respiration by transmitting the force directly to the membrane sector of the rotor via a Grotthuss chain of water molecules (Spikes et al., 2020). In mammals, this membrane sector of the rotor is composed of a ring of eight c-subunits, and is attached to an extramembranous central stalk assembly of subunits γ, δ and ε, which penetrates into the extramembranous α_3_β_3_-catalytic domain *ca*. 90 Å distant from the membrane domain (Watt et al., 2010; Spikes et al., 2020). Three of the six interfaces between α-and β-subunits contain the catalytic sites of the enzyme, and as the rotor turns in 120^°^ steps during the rotary cycle, these sites undergo a series of structural changes that are associated in turn with the binding of substrates, and the formation and release of ATP from each site (Walker, 2013). The catalytic domain is also attached to the membrane domain via a peripheral stalk (PS), an elongated rod that reaches *ca*. 150 Å between the catalytic domain and the surface of the inner mitochondrial membrane (IMM) (Collinson et al., 1994; Collinson et al., 1996; Spikes et al., 2020). The core of the rod is provided by a long α-helix, bH3, which extends between the α_3_β_3_-catalytic domain and the IMM, and then penetrates and crosses the IMM to reach the inter-membrane space and then returns to the mitochondrial matrix via a second transmembrane span. The membrane extrinsic part of bH3 is supported and largely rigidified by other α-helices in the d- and F_6_-subunits and in the membrane extrinsic region of subunit ATP8, all bound approximately parallel to the membrane extrinsic region of bH3. The upper part of the rod is attached via the C-terminal α-helix of subunit b, bH4, to the C-terminal domain of the OSCP subunit, and the N-terminal domain of this subunit binds to the N-terminal regions of the three α-subunits. The two domains of the OSCP are joined by a flexible linker, which provides the PS with a universal joint. During the rotary catalytic cycle, the α_3_β_3_-domain rocks from side to side, as noted before (Zhou et al., 2015; Sobti et al., 2019; Guo et al., 2019; Sobti et al., 2020), with little or no displacement perpendicular to the lateral motion of the PS towards the central axis of the rotor (Spikes et al., 2020), and the PS stalk accommodates this rocking motion via the combined action of the universal joint, and a hinge in the PS close to the surface of the IMM (Spikes et al., 2020). The membrane bound N-terminal region of the b-subunit is folded into two further α-helices, bH1and bH2, and together with the transmembrane domain of bH3, they form the skeleton of a wedge-shaped structure in the membrane domain (Spikes et al., 2020), with bH1 sitting on top on the matrix side of the IMM and transmembrane α-helices bH2 and bH3 subtending an angle of *ca*. 45^°^. Supernumerary subunits e, f and g also contribute to the structure of the wedge. The transmembrane α-helices eH1 and gH3 of subunits e and g augment α-helix bH2, and the top of the wedge on the matrix side of the membrane is provided by four amphipathic α-helices, two in each of the g- and f-subunits, lying in the lipid head-group region. Internal cavities in the wedge are occupied by five specifically bound lipids, three cardiolipins (CDL1, CDL2 and CDL3) and two less well defined phospholipids modelled tentatively as phosphatidyl glycerols (LHG4 and LHG5). These phospholipids enhance the stability of the wedge (Spikes et al., 2020). Subunit j is associated with the wedge, and in the dimeric complex the two j-subunits interact with each other across the monomer-monomer interface. In the inner membranes of the mitochondria, the dimeric ATP synthase complexes occupy the tips of the cristae, and the dimers are arranged in long rows along the tips of the cristae (Dudkina et al., 2006; Davies et al., 2011; Blum et al., 2019). How the dimers are held together in higher oligomers is uncertain, but protein-protein interactions in the membrane domains of dimers may provide inter-dimer tethers (Gu et al., 2019; Spikes et al., 2020). If the capacity of the monomeric complexes to form dimers is removed, for example by deletion of one of the wedge components, the structures of the inner membranes change dramatically, and the cristae tubes disappear (Arselin et al., 2004; Rabl et al., 2009; Davies et al., 2012; Habersetzer et al., 2013; Jackson et al., 2017; Oláhová et al., 2018; Siegmund et al., 2018). Thus, the dimerization of the ATP synthase appears to be a determinant of the formation of cristae. These dynamic ATP synthase complexes operate in the ever changing structural context of the mitochondria themselves. The organelles are constantly being re-modelled by fission of mitochondrial networks and by fusion of vesicular mitochondria, influenced, for example, by cellular conditions such as energetic state, the cell-cycle or by pro-apoptotic or cell-death factors (Giacomello et al., 2020). The fusion and fission events involve changes in both inner and outer membranes, in the latter case again influencing the structures of the cristae and the disposition of the dimeric ATP synthase complexes within them. During these dynamic events and under conditions of active cellular growth, the inner membranes of the mitochondria need to remain coupled to ATP synthesis. Therefore, in order to remain active, the dimeric ATP synthase not only has to accommodate changes in the monomer-monomer interface that arise directly from its own catalytic activity, but also others that stem from the dynamics of the organelle itself, and without dissipating the pmf by leakage of protons through the monomer-monomer interface. In tomographic reconstructions of mitochondrial cristae, zig-zagging, lateral bending, perpendicular curvature and non-uniform packing of rows of dimers of ATP synthase have been observed (Davies et al., 2011; Davies et al., 2012; Daum et al., 2013), illustrating the wide range of varying modes of oligomerization under which the ATP synthase complexes operate.

As shown here, the interaction between the two wedges linking the monomers in the dimeric ATP synthase is not a constant feature. It changes to accommodate both the rocking motions of the membrane extrinsic catalytic domain associated with catalysis, and other motions that are independent of catalysis observed in the isolated dimeric ATP synthase. These pivoting motions and sliding translations between the monomeric complexes are intrinsic features of the dimeric ATP synthase. They allow it to operate in the ever-changing ultrastructure of the inner membranes of mitochondria and also to contribute to their ultrastructure.

## RESULTS AND DISCUSSION

### Classification of dimeric particles of bovine ATP synthase

As described before, three data-sets of 4,267, 2,238 and 4,096 dose-fractionated exposures of the purified dimeric bovine ATP synthase were collected, and structures of the monomeric enzyme were produced by single particle analysis of 176,710 particle coordinates (Spikes et al., 2020). Complete structures of the monomeric complex were built in three rotational states named 1, 2 and 3 (Spikes et al., 2020). By combining these three states (see Table S1), models of dimeric complexes were constructed in a range of combinations of catalytic states and sub-states (see Scheme S1 and Table S2). The sub-states arise from changes that are independent of catalysis in the relative disposition of the two monomers to each other (see Scheme S2). By hierarchical classification of the earlier data-set (Scheme S1) and subsequent refinement, 154,130 particles were resolved into nine discrete classes of the dimeric assembly representing rotational states [s1:s1], [s1:s2], [s1:s3], [s2:s1], [s2:s2], [s2:s3], [s3:s1], [s3:s2], and [s3:s3] at resolutions of 9.2, 11.9, 9.0, 9.4, 8.5, 10.7, 9.7, 11.4 and 13.1 Å, respectively. Composite dimer models were created by rigid body fitting of the maps and associated atomic coordinates (Spikes et al., 2020) into these lower resolution envelopes. Further classification of the particles in each catalytic state (Scheme S2) revealed fifty-nine additional sub-states of the dimer, at resolutions ranging from 13.8-23.8 Å. The sub-states are distinguished from discrete classes of the dimeric assembly in defined rotational states by sequential lettering and are grouped according to catalytic state. For example, the first, second and third sub-states of the ATP synthase dimer in which the right and left monomers are in catalytic states 1 and 3, respectively, are referred to as dimer sub-states [s1:s3a], [s1:s3b] and [s1:s3c].

### Changes in the monomer-monomer interface during catalysis

In the dimeric complex, the monomer:monomer interface in the membrane domain consists of contacts between the two j-subunits each on the external surface of the wedge (Fig. 1 and Fig. S1 *A*). From residues 1-20, subunit j is folded into an amphipathic α-helix jH1, which lies in the lipid head group region on the matrix side of the IMM where it interacts with the C-terminal region of transmembrane α-helix A6LH1 and CDL1. The negatively charged headgroup of CDL1 is bound to jK8, and to residues fQ38 and fY42 of the f-subunit in the wedge and residues aT33, and A6LK27 and A6LK30 (see Fig. S1 *B*). Transmembrane α-helix jH2 (with residues 22-39 forming the transmembrane span) lies adjacent to A6LH1 on the opposite side of aH1 to the wedge, and is associated with subunit a via interactions between its N-terminal region and residues 107-108, 110-111 and 114-115 of aH4 (see Fig. S1 *A,B*). The C-terminal region of jH2 (residues 40-49), protrudes into the IMS. Residues j50-60 were not modelled as they were not well defined in the cryo-em reconstructions. They are predicted to have an extended structure (see Fig. S2) that interacts with the same region in the adjacent j-subunit, as indicated by the structures of the dimers in various rotational states and many of their sub-states, where density beyond the modelled residues can be observed (see Fig. S1 *D-F*). However, the interactions between the two j-subunits are mobile, and the observed angle between the central axes of the central or peripheral stalks of the monomers ranges from *ca*. 75-86^°^ in the nine rotational catalytic states (see Movies 1 <u>and</u> 2). This range of angles arises by the rigid body of the wedge and the rest of the membrane domain pivoting about contact points between the amphipathic α-helices jH1 (residues 1-20), provided by residues 3, 7, 10, 11 and 14 (see Fig. 2 and Fig. 3). The two jH1 α-helices are oriented in the plane of the IMM on the matrix side as the enzyme contorts during the rotational cycle (see Figs. 1 and 2, and Movie 3). Residues 3, 7, 10, 11 and 14 in jH1 project toward the interface possibly interacting via an intervening lipid. This pivoting reduces the net side-to-side displacement of the catalytic domain arising from the rotation of the asymmetrical central stalk, and accommodates other changes in the structure of the dynamic mitochondrial cristae.

**Fig. 1.**
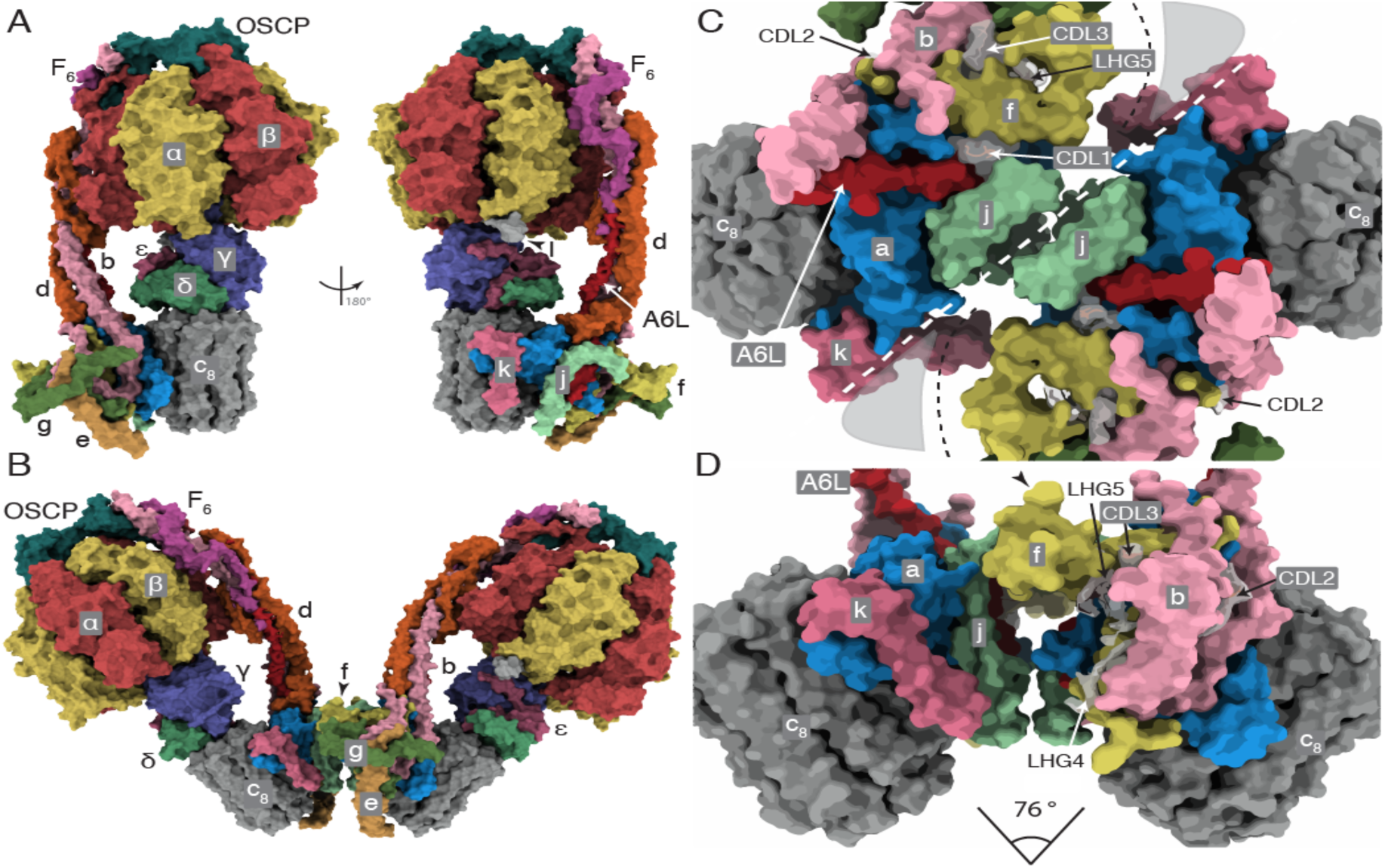
The structure of the bovine ATP synthase and the monomer-monomer interfaces in the membrane domains of dimers. *A* and *B*, the structures and subunit compositions of the bovine ATP synthase monomer in rotational state 3 and the dimer in state [s1:s1], respectively. The α-, β-, γ-, δ- and ε-subunits of the F_1_-catalytic domain are red, yellow, blue, indigo and green, respectively, with the central stalk (subunits γ, δ and ε) attached to the c_8_-ring (dark grey) in the membrane domain in contact with subunit a or ATP6 (cornflower blue). The PS subunits, OSCP, b, d and F_6_ are teal, light pink, orange and magenta, respectively, and the A6L subunit is brick red. In the region of the monomer-monomer interface, subunits e, f, g, j and k are khaki, straw yellow, forest green, sea-foam green and dark pink, respectively. Cardiolipin (CDL) and phosphatidyl-glycerol (LHG) are transparent grey. In *A*, the monomeric complex is viewed in two rotated positions, to reveal the positions of all subunits, with the rotatory axes aligned vertically. I denotes residues 1-60 of the inhibitor protein IF_1_. In *B*, the dimer is viewed from within the plane of the IMM. *C* and *D*, the solvent excluded molecular surfaces of subunits in the membrane domains of the dimeric complexes viewed between the two peripheral stalks; in *C*, the complex is viewed from the matrix side of the IMM with the monomer:monomer interfaces indicated by the white dashed line. The black dashed line denotes the protein boundary between the monomers, adjacent to a region occupied by non-specific lipids and the detergent micelle (grey shading); *D*, the orthologous view in the plane of the IMM with the monomer:monomer interfaces exposed by removal of subunits e, g and d. The angles between the axis of rotation in each monomer indicated beneath were estimated by calculating two centroids for residues 2 and 38 in each bovine c_8_-ring. The axis connecting the two centroids approximates to the rotatory axis of the c-ring. The angle of intersection was measured from the models aligned with these axes orthogonal to the direction of the view.

**Fig. 2.**
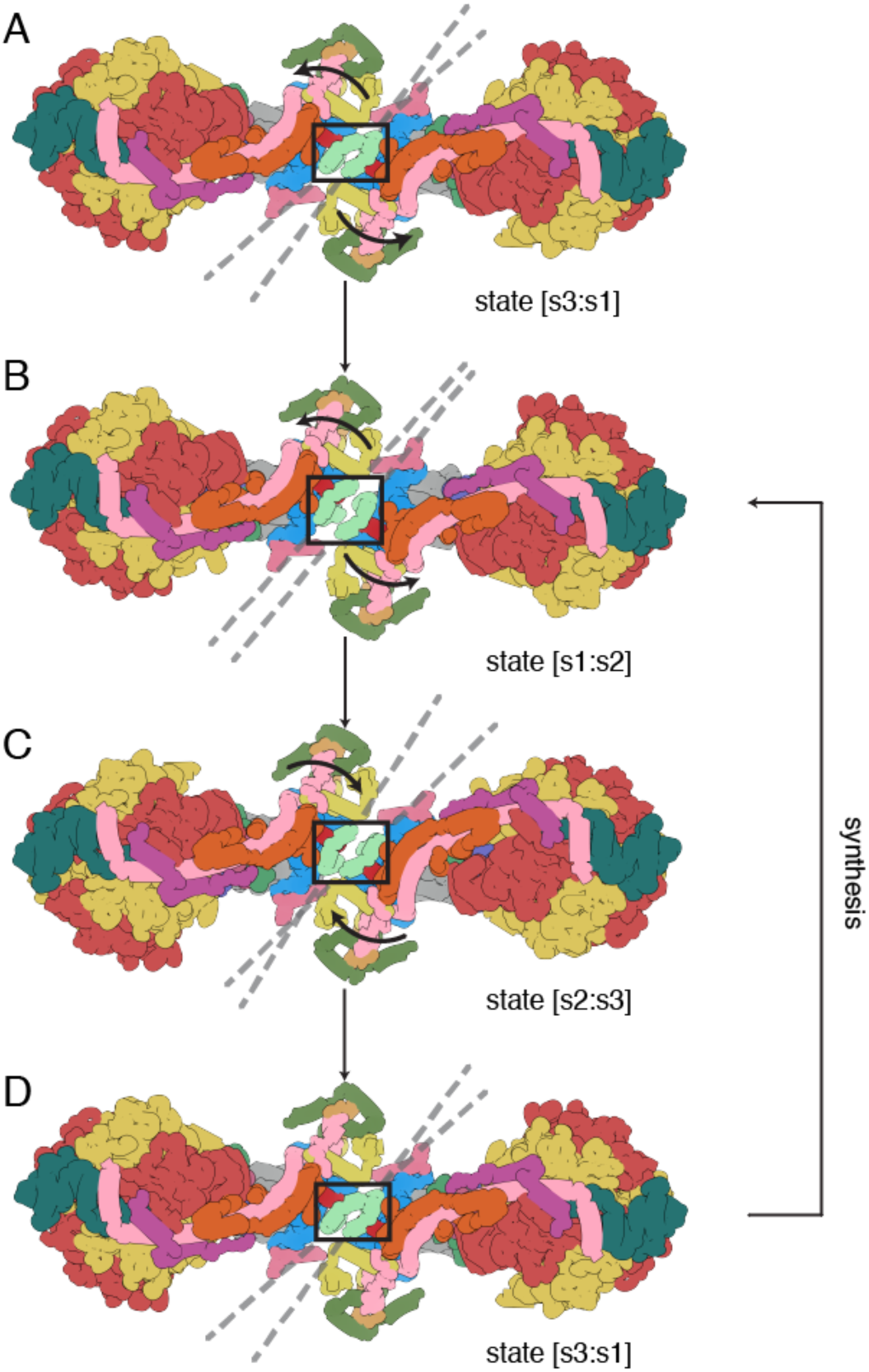
Pivoting of the membrane domains of bovine ATP synthase about the matrix contact between j-subunits during ATP synthesis. Subunit j is sea-foam green inside the black box. For colors of other subunits, see the legend to Fig. 1. *A*, the state [s3:s1] dimer. Dashed lines indicate the central axis of amphipathic α-helices jH1 (residues 1-19), which lie in the plane of the matrix leaflet of the membrane. During the synthetic rotary cycle, each monomer progresses from state 1, to state 2, to state 3 and so forth. *B*, rotation of the membrane domain about the contact point in jH1 during the transition from state [s3:s1] to state [s1:s2], with accompanying displacement of subunits e, f and g, and the transmembrane α-helices bH2 (residues 33-47) and bH3 (residues 55-73), moving the wedge outwards or inwards as indicated by the arrows. The membrane extrinsic F_1_-domains remain approximately stationary, and the rocking motion associated with the asymmetry of the central stalk is transmitted to the membrane domain via the universal joint between N- and C-terminal domains of the OSCP in the PS (ref); *C*, a similar pivoting motion occurs in the transition from state [s1:s2] to state [s2:s3]; *D*, completion of the rotary cycle by the transition from state [s3:s1] to state [s1:s2]. The synthetic rotary cycle continues via *B* as indicated by the arrow on the right. The scheme was constructed with composite atomic models of monomeric bovine ATP synthase, which were rigid body fitted into the consensus dimer structures produced according to Scheme S1. This Figure relates to Movie 3. In both this Figure and Movie 3, side chains have been removed and secondary structure elements have been dilated to produce the diagrammatic nature of the figure, which is inferred from lower resolution structures of the intact dimeric complex.

**Fig. 3.**
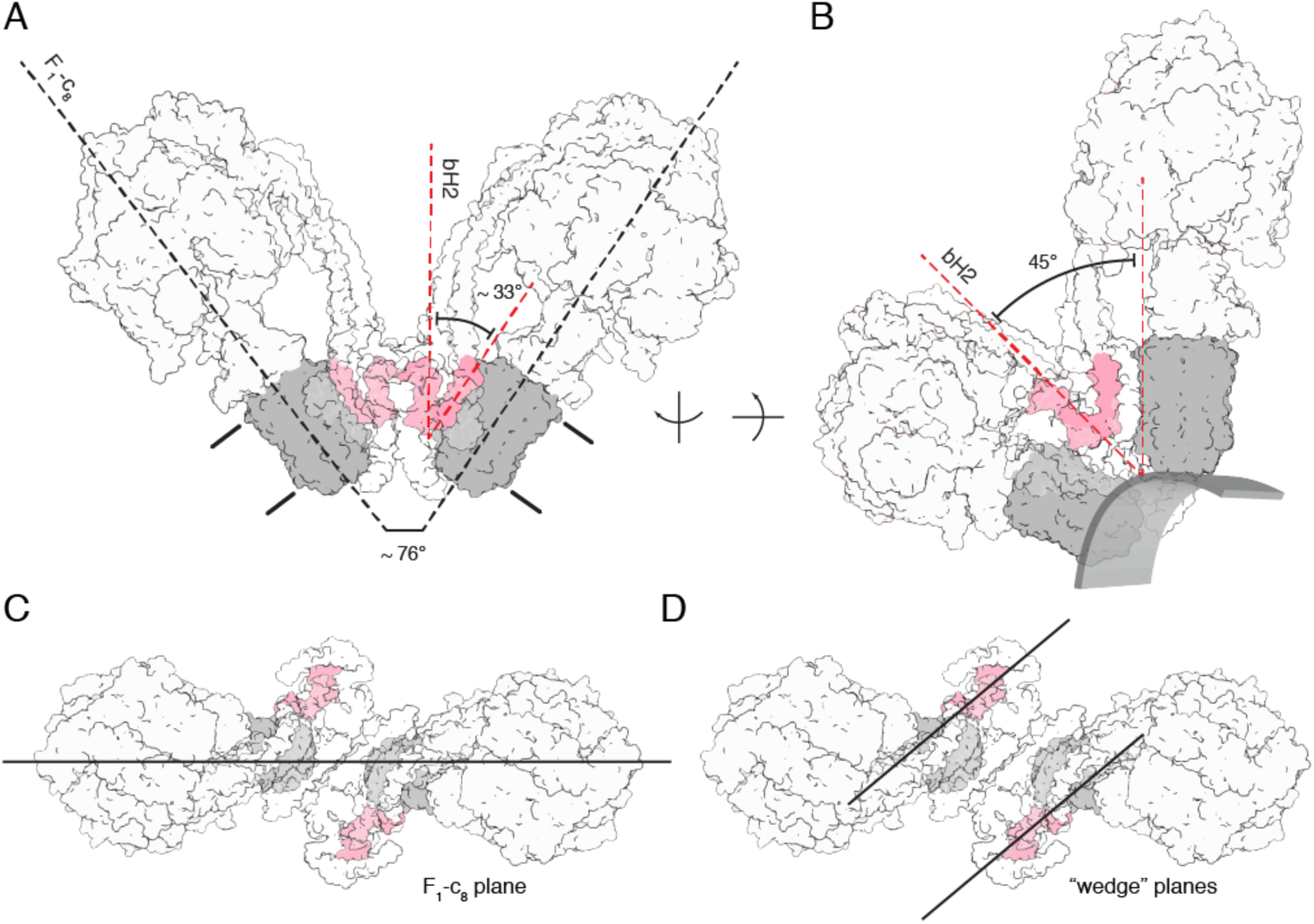
Relationship between the inclined membrane wedge in the monomeric membrane domain and the association of two monomers into a dimer. The dimer in state [s2:s2] is shown. The c_8_-ring is grey, and transmembrane α-helices bH2 and bH3 of subunit b (pink) define the wedge angle (Spikes et al., 2020). The rest of the structure is shown in silhouette. *A*, side view showing the relationship between the angles of rotatory axes and the wedge angles when measured from the same reference point, with both rotatory axes aligned in plane. The angle between bH2 and bH3 in this view is 33^°^, approximately half of the dimer angle. The approximate boundary of the membrane is indicated; in *B*, the rotatory axis is aligned vertically and the structure is rotated so that bH2 and bH3 in the wedge of the right-hand monomer are in the plane of the paper. The 45^°^ wedge angle is a measure of the inclination of the transmembrane α-helix bH2 relative to bH3 and the rotatory axis of the enzyme, as required for the wedge to provide the membrane curvature in the membrane domain of each monomer. The approximate boundary of the lower leaflet is indicated to provide perspective; *C* and *D*, top views showing the differences between the angle measurement planes to explain why a simple doubling of the wedge angle does not produce the rotatory axis angle. In the context of the angle variations observed in the consensus reconstructions, the wedge angle and dimer, or rotatory axis, angle are separate considerations. The wedge angle does not change, rather, the way in which the membrane domains are arranged relative to one another does, as described in Fig. 2 and Movie 3.

### Changes in the monomer-monomer interface unrelated to catalysis

Other variations in motion occurring within groups of particles of a defined catalytic state are sub-states that are independent of catalysis. They fall into two categories, namely those where the angle between rotatory axes is less than 90^°^ (Movie 4) and others where the angle is greater than 90^°^ (Movies 5 and 6). The motions in the former category are similar to those that occur during catalysis in that the relative dispositions of the membrane domains change by a pivoting motion about the interface between j-subunits, with the C-terminal IMS protrusions remaining in contact. The flexibility of this pivot is an intrinsic property of the interface and it has other roles beyond that described above of dissipating the rocking of the catalytic domain in the mitochondrial cristae. Most notably, it provides a mechanism to accommodate the general fluidity of the membrane in the region of curvature associated with the dimeric ATP synthases and the rearrangements that occur. The second category of motions between sub-states, where the angle between rotatory axes exceeds 90^°^, arises from sliding and twisting along the monomer-monomer interface presented by the a and j subunits and the membrane wedge, thereby effecting a compound rotation of one monomer relative to the other (see Fig. 4 and Movies 4 and 5). For example, in Fig. 4, where the structures of the ATP synthase dimers in the sub-states [s2:s2c] and [s2:s2b] are compared, the positions of the protrusion of subunit j into the IMS and of the loop region (residues 12-19) in subunit a differ significantly in the two structures (see Movie 5 and 6). In sub-state [s2:s2c], the C-terminal residues of subunit j are in contact in the intermembrane space (IMS) as in Fig. 4*C*. In contrast, in sub-state [s2:s2b], these residues are displaced by *ca*. 20 Å and the two loops consisting of residues 12-19 of subunit a appear to make contact with each other, as in Fig. 4*D*. This compound rigid body rotation of the monomers changes the angle between the rotatory axes significantly, and also re-arranges the amphipathic α-helices lying in the IMS leaflet of the membrane (see Fig. 4 and Movies 5 and 6). In the mitochondrial inner membranes themselves, this transformation could be part of a mechanism for the dimers and oligomeric rows to adapt to the convolutions of the cristae, with the two contact positions representing a simple steric mechanism that restrains the rotation to upper and lower limits. In this mechanism, the wide angle conformation (as observed in sub-state [s2:s2b]; see Fig. 4*B, D* and *F*) rests at the position where the two a-subunits are in contact and is prevented from widening further by aH1, and the conformation with a shallow angle (for example, the sub-state [s2:s2c] in Fig. 4*A, C* and *E*) rests at the position where the two j-subunits are in contact in the IMS. There is probably a continuum of sub-states between the two extremes. The conformers with shallow angles between rotatory axes of *ca*. 76^°^ to 86^°^ were the most abundant observed sub-states (see Fig. S3 and Scheme 2). To demonstrate the highly dynamic nature of the dimeric complex, all the observed conformers are compiled in Movie 7.

**Fig. 4.**
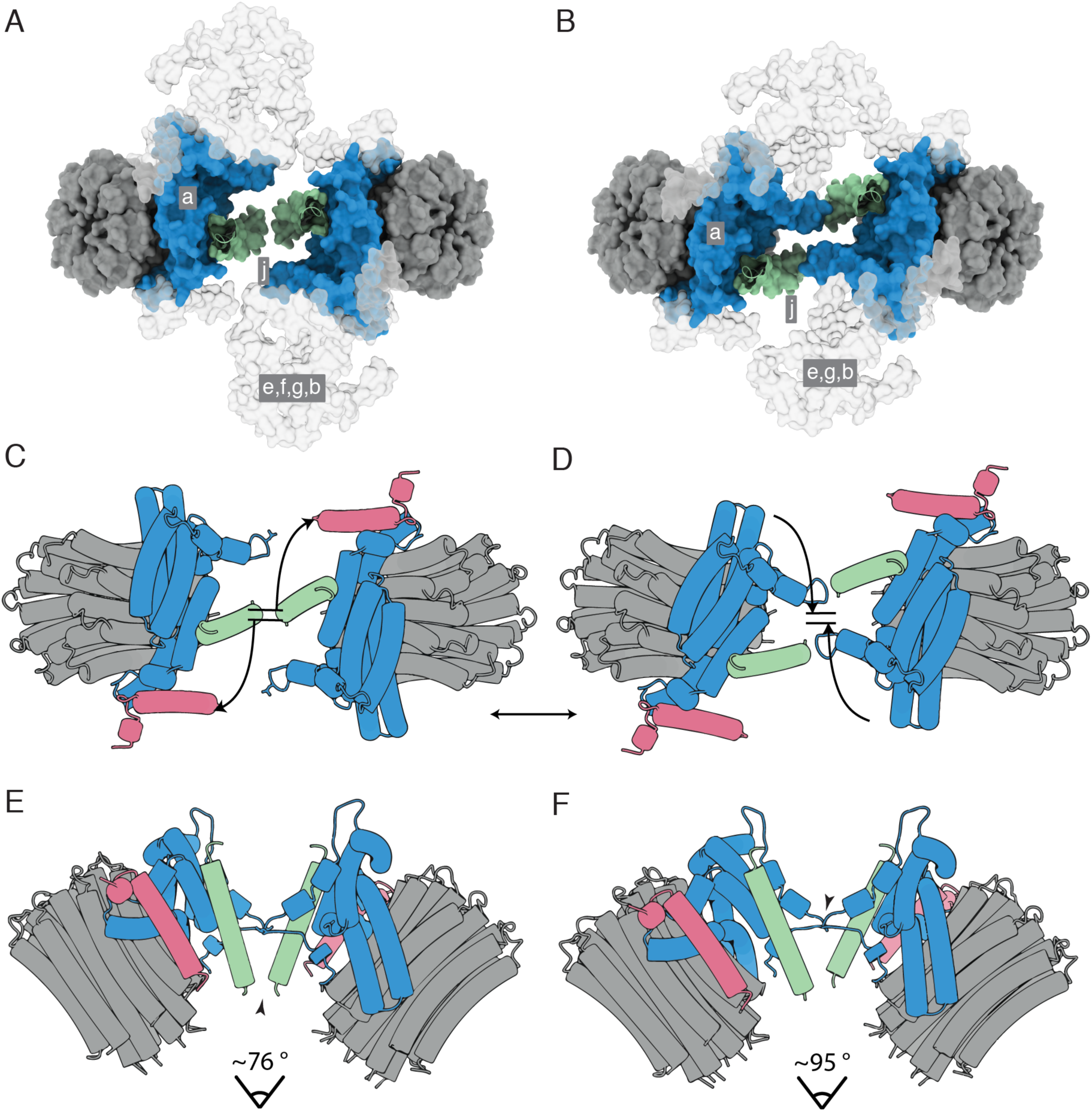
Dispositions of subunits a and j in the monomer-monomer interface of dimeric bovine ATP synthase at narrowest and widest extremities of angles between rotatory axes. Subunits a, j and k are blue, green and pink, respectively, and the c_8_-rings are grey. *A* and *B*, views from the matrix side of the IMM of the monomer-monomer interfaces in the membrane domains of bovine dimers with angles between rotatory axes of *ca* 76^°^ and *ca* 95^°^, respectively. In *A* and *B*, respectively, the dimeric sub-state [s2:s2c] (see Schemes 1 and 2), and *B* in the dimeric sub-state [s2:s2b] (see Scheme 2) are shown as examples. The positions of subunits e, f, g and the membrane domain of subunit b are indicated in grey silhouette. For clarity, the F_1_-domains and all PS subunits have been removed. *C-F*, schematic representation of the monomer-monomer interfaces, viewed in *C* and *D* as in *A* and *B*, respectively, and in *E* and *F* orthogonal to the plane of the membrane. In *E*, the arrowhead indicates an interaction between the C-terminal residues of the two j-subunits. In *F*, the arrowhead indicates a possible interaction of residues 12-19 in a loop proceeding αH1 of the a-subunit. In *C* and *D*, the curved arrows indicate the approximate compound rotation of the membrane domains that occurs as the angle of the rotatory axis changes between the dimer sub-states. This Figure relates to Movies 5 and 6.

It cannot be excluded that the wide angle sub-states with rotatory axis angles greater than *ca*. 95^°^, which depart significantly from the consensus structures with rotatory axis angles of *ca*. 76-86^°^, arise from the influence of the detergent micelle and the lack of the interactions between dimers that occur in native membranes. The sub-tomographic reconstructions of dimeric ATP synthases from mammalian mitochondria, where rotatory axis angles of *ca*. 80 ^°^ have been reported, represent the average structures of many individual molecules, or sub-tomograms, within the data (Davies et al., 2011), similar to the consensus dimer in Scheme S1. Therefore, they do not exclude the possibility of large variations in the angle of the rotatory axis, especially if those conformations arise rarely, for example at cristae ultrastructures with sharp negative curvature, or during dynamic rearrangements of the cristae, and indeed the dimer sub-states with wider angles are relatively rare (see Fig. S3*C* and *D* and Scheme S2). Although it is not certain whether these very wide rotatory axis angle substates occur in a native membranes, as the analysis was performed on isolated dimers, the data strongly suggest that the interfaces between monomers are significantly dynamic and that this dynamism is a general property of the interface, linked to motions of the enzyme that are both dependent and independent of catalysis. Indeed this dynamism, which manifests as pseudo-c2 symmetry about the dimer interface, necessitated specific refinement strategies to prevent the rotation of particle orientations about the pseudo-c2-axis in order to isolate and subsequently maintain catalytically homogenous particle sub-sets during further particle-realignment (see Scheme 1). Therefore, during image processing, each monomer was given the arbitrary designation “left or “right” with respect to a single viewing direction, and, for example, the state [s1:s2] and state [s2:s1] dimers are distinct and do not represent the same molecular state.

### Interactions between dimers

The issue of how dimeric ATP synthases interact with each other in the long rows that form along the apices of the cristae in the IMM cannot be resolved definitively by the current study of the structure of the isolated bovine dimers. However, the structures and other biochemical information suggest possibilities of how the dimeric ATP synthases might form tetramers and higher oligomers involving homo-interactions between the two copies of each of subunits g and k. In the dimeric bovine structure, the 102 amino acid residue g-subunit is folded into three α-helices, gH1 (residues 20-36), gH2 (residues 42-60) and gH3 (residues 69-93) with transmembranous gH3 augmenting the skeleton of wedge, and amphipathic gH1 and gH2 associated with the top of wedge on the matrix side of the IMM. In the dimeric complex, the two gH2s have the capability to contact each other across the dimer-dimer interface and link the dimers together, as depicted in Fig. 5, with weak interactions across the dimer-dimer interface between residues gQ50, gK53 and gK54 (see Fig. 7) in each protomer providing a potential molecular basis for the requisite sliding, translational and rotational elements to generate the wide range of cristae structures that have been observed in mitochondria. Such a fluid interface would allow, for example, the long rows of ATP synthase oligomers to follow the curvature along the apices of the cristae (see Fig. 5*B, C* and *D*), and to readjust their positions as the curvature changes, without impeding the dynamics of the cristae. This fluidity is illustrated in Fig. 5*B*, where the ATP synthase dimer follows the curvature of the inner membrane along the axis shown in Fig. 5*C* and 5*D*. The fluidity of the dimer-dimer interface could allow either a rotation about a fixed contact point between g-subunits, or dislocate the contact and translate the dimers sideways (see Fig. 6*A*). These adjustments of inter-dimer contacts, and combinations of them can engender a range of oligomeric arrangements. They include non-uniform packing (Fig. 6*B*-*G*), where rotation about a fixed contact point produces lateral bends in oligomer rows (as viewed from above; see Fig. 6*F* and 6*G*), and where contact dislocation permits the dimers to stack in a compact arrangement (Fig. 6*C* and *E*), or to form staggered rows (Fig. 6*D*). These arrangements are constrained geometrically in the cristae by the spherical catalytic domains of the enzyme. Thinning of the footprint of the membrane domain of the dimer, as viewed from above or below the membrane, resulting from changes in the angle between the rotatory axes as described in Fig. 4 and in Movies 5 and 6, places adjacent catalytic domains in closer proximity, and therefore limits the degree of lateral curvature of the oligomeric row. Such a mobile contact also accounts for the positive and negative curvature along the cristae rows (see Fig. 5*C* and *D*), and, in a variety of combinations with dislocations, produces zig-zagging, lateral bending, perpendicular curvature and non-uniform packing. The oligomer rows may also deform plastically, much like an armature wire in a sculpture model, providing strength to maintain the apex of the cristae over long distances and simultaneously allowing different types of ultrastructure, such as lateral and perpendicular bends, to form. Many of these arrangements have been observed in tomographic reconstructions of the cristae (Davies et al., 2011; Davies et al., 2012; Daum et al., 2013).

**Fig. 5.**
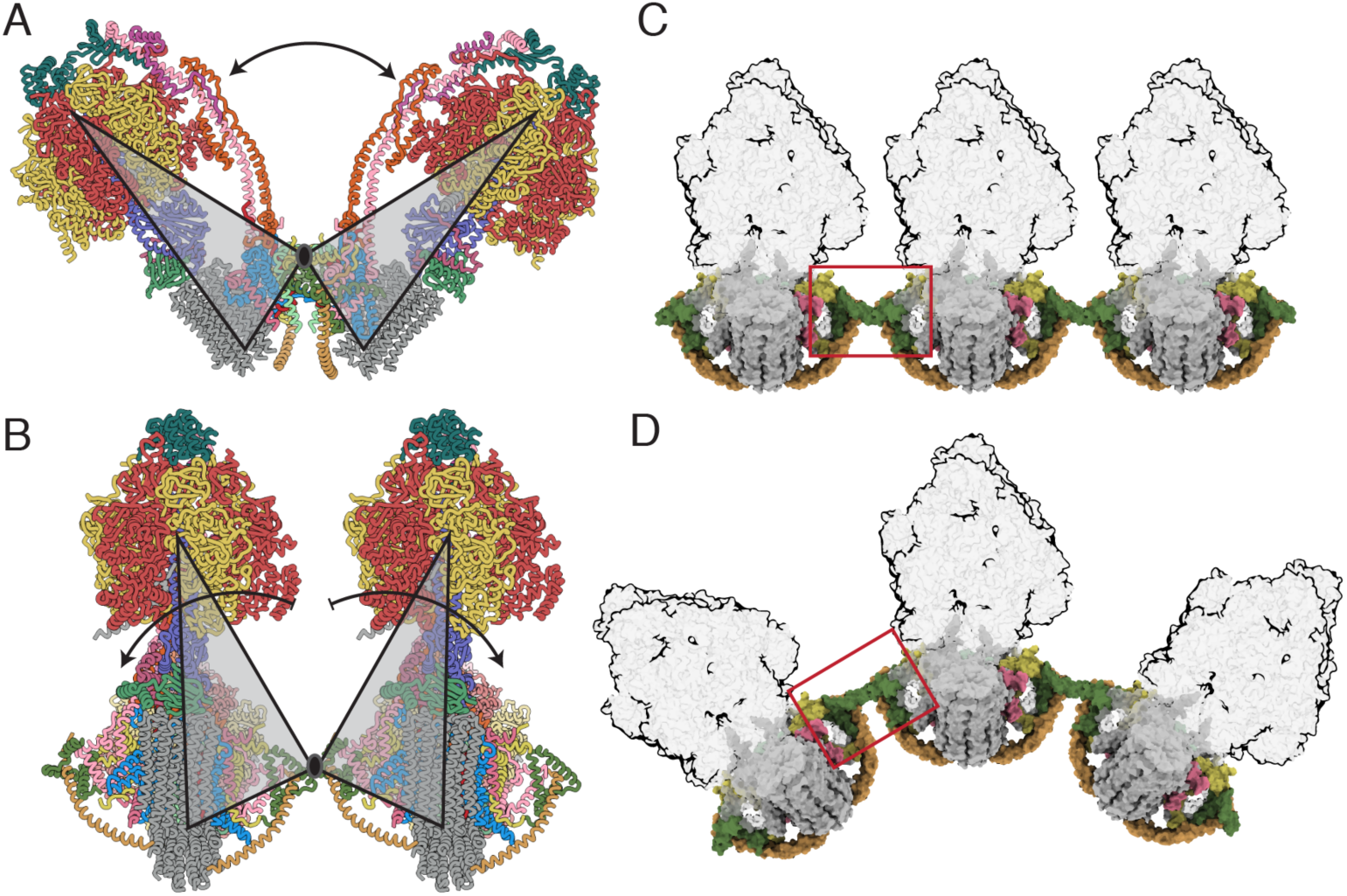
Accommodation of the dimeric ATP synthase to the membrane curvature in mitochondrial cristae along the axis of oligomerization via flexible inter-dimer contacts. *A*, a view, orthogonal to the plane of the IMM, of the bovine ATP synthase dimer in state 1: state 1. Grey triangles represent the monomers, moving back and forth as rigid bodies by rotation about their point of contact in j-subunits; the rotatory axes of both monomers are aligned in the same plane, and the arrow indicates the range of angles between rotatory axes; *B*, side view, parallel to the plane of the IMM, of two monomers in a tetrameric arrangement of two adjacent dimers formed via contacts between g-subunits. The arrows indicate the independent courses of their catalytic domains by rotation about this point; *C* and *D*, similar view to *B*, with the F_1_-domains in silhouette illustrating how changes in the interface between dimers (red box) allow a curved ultra-structure to develop along the apices of the cristae. For colors of subunits, see the legend to Fig. 1.

**Fig. 6.**
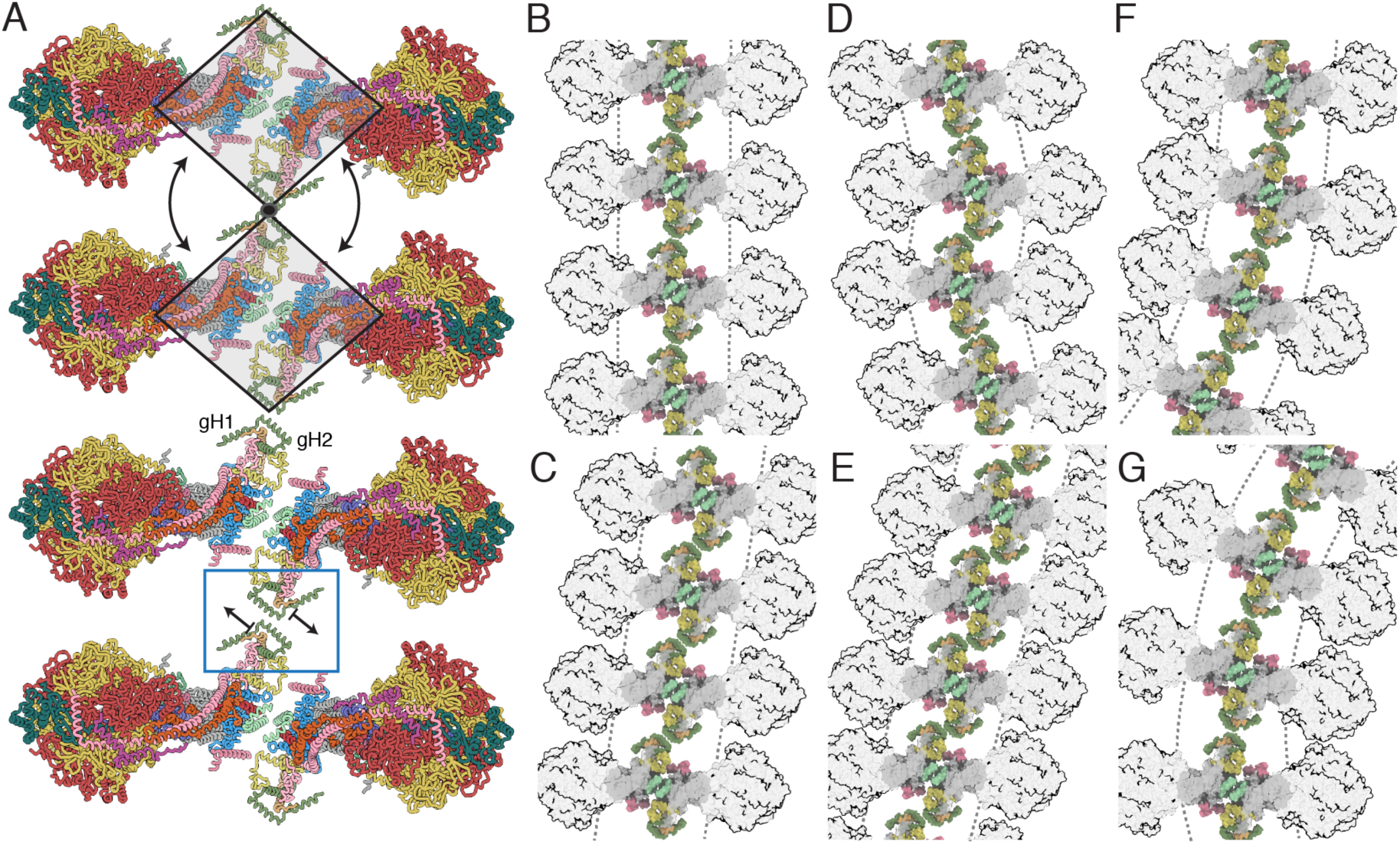
Possible modes of association of dimeric bovine ATP synthases in the cristae. *A*, row of dimers viewed from above the cristae tips. In the two upper complexes, the grey trapezoids represent individual dimers moving as rigid bodies; in the lower two complexes, the individual dimers are translated as indicated by the arrows via the dimer-dimer interface in the blue box; *B-G*, the impact of translation or rotation of dimers on the long range order of oligomeric rows. In *B*, back-to-face stacked rows of dimers provide the simplest architecture of straight and planar oligomers. *C, D* and *E*, effects of dislocation of the g-g interaction at the interface between two dimers. In *C* and *E*, compact back-to-face rows of dimers to form with a range of inclinations with respect to the perpendicular axis, or, in *D*, as an alternating “zig-zag” arrangement. In *F* and *G*, bending of the rows of oligomers laterally by rotation about a fixed point at the g-g contact. More complex arrangements can be envisaged by combination of the effects in *B-G*, with the additional possibility of curving the membrane positively along the axis of oligomerization, as in Fig. 5*D*. For colors of subunits, see the legend to Fig. 1.

**Fig. 7.**
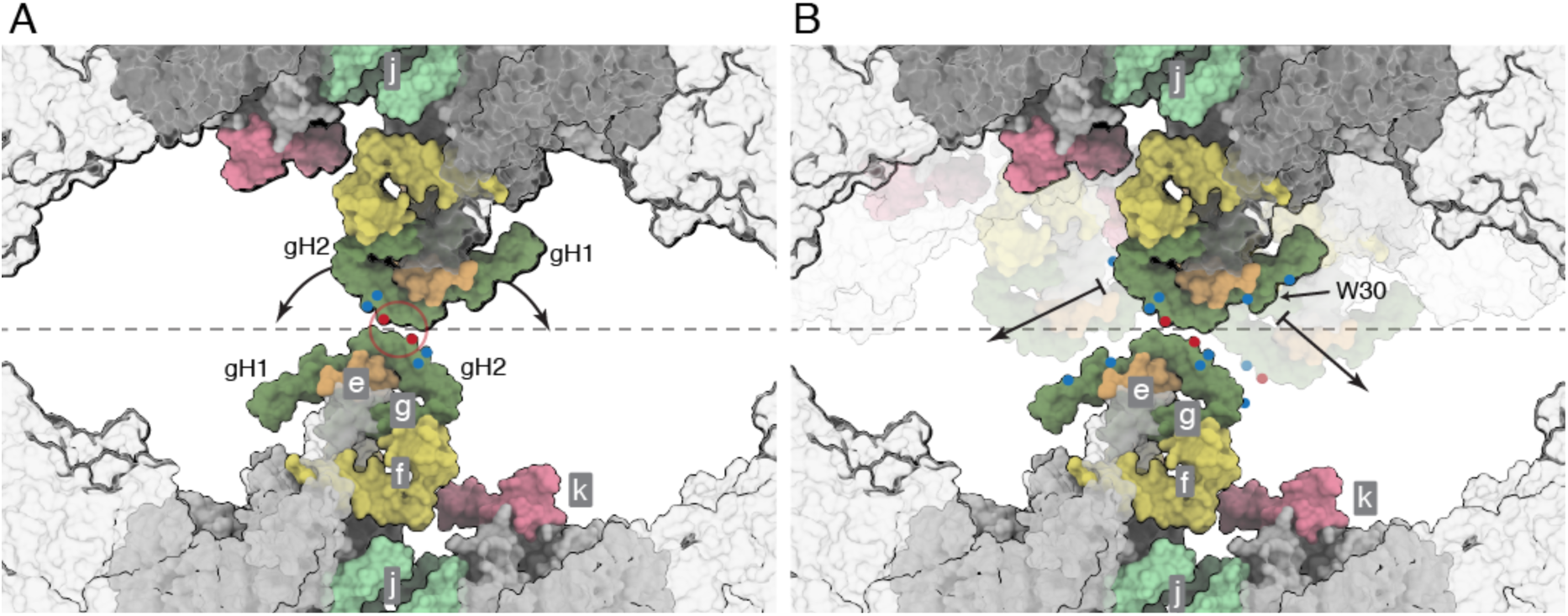
Close up of the dimer-dimer interface in a possible tetrameric bovine ATP synthase. The interactions shown as a molecular surface between g-subunits in two adjacent dimers viewed from inside the mitochondrial matrix between the axes of the peripheral stalks, as in Fig. 6*B*-*G*. Portions of the catalytic domains, situated above the viewing plane, are shown in grey transparency. The grey dashed line indicates the approximate tetramer interface, which is surrounded by lipids. *A*, rotation of the dimers relative to one another, as indicated by the arrows, about a contact point between the two g-subunits, highlighted in the red circle. Key negative and positive polar residues are indicated by blue and red dots, respectively; *B*, contact dislocation of the interface between g-subunits as indicated by the arrows. Possible positions of the translocated regions are shown in transparent underlay. Residue W30 might be involved in a hydrophobic “knobs-into-holes” interaction with regions of gH1 that are rich in leucine and isoleucine residues. Additional polar residues in gH1 and gH2 are indicated by colored dots. Subunits e, f, g, j and k are khaki, straw yellow, forest green, sea-foam green and pink, respectively. The c_8_-ring is grey.

It is also possible that subunit k could be involved in linking dimers together. During the process of assembly of the closely related human enzyme, this subunit is the last to be incorporated into complex, and in human cells grown under glycolytic conditions, subunit k turned over more rapidly than the other subunits of the enzyme which turned over together at a slower rate (He et al., 2018). These observations led to the suggestion that subunit k might participate in a mechanism for remodelling IMMs (He et al., 2018). The partial structure of bovine k is compatible with this proposal. In the dimeric bovine complex, residues 12-47 of the 57 residue subunit k were resolved (Spikes et al., 2020). The N-terminal region of the k-subunit is folded into a short α-helix, kH1, from residues 14-17 and is linked to kH2 by a loop (residues 18-23). This region is followed by the single transmembrane α-helix, kH2, from residues 24-44 which is associated with subunit a via interactions with aH4 and aH5. The C-terminal region (residues 45-57) of subunit k is in the IMS and is predicted to be extended. Thus, it is possible that in the mitochondrial membrane, this extended region could be in contact with the C-terminal region of subunit g across the dimer-dimer interface. The distance between k and g subunits in adjacent dimers decreases as the angle between rotatory axes in the dimer increases. Rearrangement of wedge subunits along the dimer interface, concomitant with changes to the rotatory axes as shown in Movies 5 and 6, narrows the membrane footprint of the dimer thereby bringing adjacent dimers closer.

The structure of the tetrameric porcine ATP synthase (Gu et al., 2019) represents another mode of interaction of two ATP synthase dimers where two IF_1_ dimers span between their globular catalytic F_1_-domains, and in addition there are contacts between the membrane domains, although the subunits involved were mis-identified (Spikes et al., 2020). In a corrected structure of the porcine tetramer, the g-subunits from adjacent dimers are in contact on top of the matrix side of the membrane via gH2 and the preceding loop (residues 39-53), the membrane domains of the e-subunits are in close contact, and residues 13 and 14 of subunit k interact with gH1 on the matrix side of the membrane (Spikes et al., 2020). In addition, its unresolved C-terminus could interact with subunits g and a on the IMS side. However, the physiological relevance of this inactive state is unclear, and the *in silico* association of tetrameric complexes leads to a closed loop within six dimeric units (Fig. S4). Thus, this conformation is unlikely to be relevant in consideration of the wider plasticity of the IMM.

The elucidation of the molecular basis of the interactions of dimers of ATP synthase in the mitochondrial cristae requires further study by high resolution tomography of the mitochondrial membranes. The current study illustrates that the intrinsic properties of the ATP synthase dimer units provide access to a wide range of topographies of the cristae.

## ACKNOWLEDGEMENTS

This work was supported by Medical Research Council (MRC) UK Grants MC_U105663150, MR/M009858/1, MC_U105663150, and MC_UU_00015/8 (to J.E.W.). From 2013 to 2017, T.E.S. was in receipt of an MRC PhD Studentship. The cryo-electron microscopes at the University of Cambridge Department of Biochemistry were funded by Wellcome Trust grants 202905/Z/16/Z and 206171/Z/17/Z (J.E.W. co-applicant/stakeholder). We thank D. Y. Chirgadze and colleagues (Department of Biochemistry, University of Cambridge) and Y. Chaban and staff of Electron Bio-Imaging Centre (eBIC), Diamond Light Source, for expert operation of microscopes and advice on data collection; and E. R. S. Kunji and M. Wikström for their helpful comments on the manuscript.

## MOVIES

**Movie 1.mp4**

**Movie 1. Structural heterogeneity in dimeric bovine ATP synthase associated with catalysis**. The 13 sec movie demonstrates the rocking motion of the catalytic domains as they step through the three main catalytic states and the corresponding contortions of the PS as it responds to the torque of rotation, and also the subtle changes in the spatial relationship of each monomer to the other as the enzyme cycles through catalytic states and how this changes the angle of the rotatory axis. In Movie 2, these subtle changes can be seen also by tracking the movement of subunit e (khaki) and subunit g (forest green). The range of observed angles between rotatory axes in the consensus reconstructions was *ca*. 76^°^ to 86^°^. In the membrane, it is likely that these movements will be damped by the lateral pressure of the bilayer and by interactions with adjacent dimers. This damping would result in larger apparent motions of the PS and catalytic domain (which can be demonstrated by aligning monomers of state 1, state 2 and state 3 via their a-subunits, that is, by fixing the relative motion of the membrane domains). Conversely, this fluidity in the membrane domain may provide a mechanism to dampen motions caused by asymmetric rotation by reducing the net displacement of the catalytic domain.

**Movie 2.mp4**

**Movie 2. The rotary cycle during synthesis and hydrolysis**. In the synthetic direction, reconstructions appear in the order state [s3:s1] > [s1:s2] > [s2:s3], and the rotor turns anticlockwise as viewed from the mitochondrial matrix; during hydrolysis with clockwise rotation, the order is reversed. This 21 sec movie complements Movie 1 by identifying the subunits of the enzyme and by displaying the reconstructions in the physiological order of catalysis. From 00:00-00:06 sec, the reconstructions are shown slowly in order, from 00:06 sec to 00:10 sec, the enzyme is undergoing synthesis, and from 00:10 sec to 00:14 sec hydrolysis. Then the sequence from 00:06 sec to 00:14 sec is repeated. Although the enzyme is inhibited by IF_1_, it is possible to construct the rotary mechanism because all three states of the catalytic β-subunits, β_E_, β_DP_ and β_TP_, are present in the structure. In the intact monomer, the three-fold rotational symmetry of the mechanism is broken by the association of the PS, and the three IF_1_ inhibited rotational states represent the full catalytic cycle in any of the three catalytic sites. As the maps or models appear in sequence, each catalytic site cycles through the β_E_, β_DP_ and β_TP_ states during synthesis, and the reverse sense during hydrolysis. As shown in the movie, the dimeric enzyme contorts significantly during the independent rotary cycles of each monomer. The α, β, γ, δ, ε, OSCP and F_6_ subunits of the membrane extrinsic domain are dull red, golden yellow, purple-blue, green, dark purple, teal and magenta, respectively. The inhibitor protein, IF_1_, (grey) protrudes from the α_DP_-β_DP_ interface in various positions. The membrane domain subunits a, A6L, b, e, f, g, j and k are cornflower blue, brick red, light pink, khaki, straw yellow, forest green, sea-foam green and dark pink, respectively. The phosphates of the lipid headgroups are bright red. The c_8_-ring and remaining unmodelled density of the micelle are dark grey and light grey, respectively.

**Movie 3.mp4**

**Movie 3. Pivoting of the membrane domains of adjacent monomers of bovine ATP synthase about the matrix contact between j-subunits during catalysis**. The 53 sec movie, starts with the consensus reconstruction of the ATP synthase dimer in state [s1:s2] (grey) followed by a schematic representation of the monomeric models rigid body fitted into the density. Then from 00:05 sec to 00:013 sec, the state [s2:s3] and state [s3:s1] dimers and their atomic models appear in catalytic order during synthesis. From 00:14 sec to 00:30 sec, side and top views of the enzyme highlight the rearrangement of the membrane domains during synthesis. In particular, there is significant movement in the relative positions of subunits f (straw yellow) and g (forest green), which rotate with the rest of the membrane domain during the pivoting motion. Finally, transitions between pairs of catalytic states are alternated to accentuate each component of the total movement. From 00:30 sec to 00:37 sec are shown the transition from state [s3:s1] to state [s1:s2]; followed by the transition from state [s1:s2] to state [s2:s3] from 00:38 sec to 00:45 sec, and the transition from state [s2:s3] to state [s3:s1] from 00:45 sec to 00:53 sec; in the top view, the changes at the interface between the two j-subunits can be seen, about which the monomers pivot during the catalytic cycle. For colors of subunits and other details, see the legend to Fig. 1.

**Movie 4.mp4**

**Movie 4. The fluidity, independent of catalysis, of the monomer:monomer interface at rotatory axis angles less than or equal to 90**^**°**^. The 56 sec movie depicts a transition of the state [s2:s2] sub-states from sub-state [s2:s2c] > [s2:s2f] > [s2:s2g] > [s2:s2c]. The sub-states were derived by classification of ATP synthase dimer particles in rotational state [s2:s2]. Thus, the pivoting is independent of both the action of catalysis and the associated motions described in Movies 1 and 2, although the movement mode is similar. The transitions between these sub-states demonstrate the pivoting motion about the subunit j:subunit j interface in the centre of the membrane domain, with the accompanying rearrangement of subunits e, f, g and k. In the first part of the movie (00:00 sec to 00.21 sec) a side view, in the plane of the membrane, is presented, and in the second part, from 00:22 sec to 00:40 sec, the same sub-state transitions are viewed from the mitochondrial matrix. From 00:40 sec to 00:56 sec, the view is from the inter-membrane space showing the C-terminal contact point of subunit j protruding into the inter-membrane space. Several sub-states displaying this feature were found in all consensus dimer particle sub-sets and, when combined, they represent the most abundant structural conformers in the dataset with 67.6 % of particles belonging to classes in which the reconstruction displays evidence of contact between the C-terminal regions of j-subunits. In general, the foot-print of the membrane domain of these sub-states is wider, and the angle between the central axes of the F_1_-c_8_ domains is narrower with a correspondingly shorter distance between F_1_ domains. The membrane domain subunits a, A6L, b, e, f, g, j and k are cornflower blue, brick red, light pink, khaki, straw yellow, forest green, sea-foam green and dark pink, respectively. The c_8_-ring and remaining unmodelled density of the micelle are dark grey and light grey, respectively.

**Movie 5.mp4**

**Movie 5. Trajectory, independent of catalysis, towards the formation of a wide angle between the central axes of the F**_**1**_**-c**_**8**_ **domains in a dimer of ATP synthase**. The widening arises from a sliding and twisting of the membrane domains along the monomer-monomer interface surface provided by subunits a and j and occurs independently from the action of catalysis. The 36 sec movie displays transitions between several sub-states with more obtuse rotatory axis angles than those in the consensus reconstructions (shown in Movies 1, 2 and 3) and in the other sub-states of the state [s2:s2] dimer (see Movie 4). From 00:00 sec to 00:06 sec, the density of the dimer reconstructions is colored to match the subunit colors of the composite atomic models fitted within them (see Fig. 1). From 00:07 sec to 00:15 sec, the densities are interpolated in the following order; sub-states [s2:s2c], [s2:s2a], [s2:s2e], [s2:s2b], [s2:s 2e], then [s2:s2c] and so forth. This sequence is repeated in a view from the matrix and IMS sides of the complex from 00:19 sec 00:24 sec and 00:31 sec to 00:36 sec, respectively. The dramatic changes in the angle of the rotatory axes and the substantial rearrangement of the dimer interface thins the foot-print of the membrane domain significantly (as viewed from above or below the complex). This thinning affects how dimers of dimers and higher oligomers might form, as in this configuration subunit k is positioned closer to subunit g in an adjacent dimer. Also, the IMS protrusion of subunit j is absent, possibly because either a contact between the C-terminal regions of the subunit has been broken, or because of the lower resolution of these reconstructions. Other possible explanations are that this wide angle conformation results from the loss of one or all of the following: subunit j, CDL1, or unresolved lipids in the wedge. It is also possible that the extended structure of the C-terminal region of subunit j remains in contact, but was not resolved. This is possible, if not probable, as the distance between the last modelled residues in the subunit j protomers is *ca*. 29-38 Å at its furthest in sub-state [s2:s2b] and the C-terminal 11 residues of subunit j were not built. If the unresolved region comprises an extended and/or a partially α-helical structure, 22 residues would be sufficient to span the distance between their C-termini.

**Movie 6.mp4**

**Movie 6. Detailed view of the interface rearrangement, independent of catalysis, arising in the trajectory towards the wide angle dimeric ATP synthase**. A top view of the trajectory in Movie 5 is shown. Subunits a and j are cornflower blue and sea-foam green, respectively, and are shown as a solvent excluded surface. The 41 sec movie shows the sliding and twisting of each monomer along the monomer-monomer interface that leads to the widest of the rotatory axis angles, and the narrowest membrane domain footprint. The final part of the movie (from 31 sec) depicts the same sequence of events in a side view of the dimeric complex, shown as a solvent excluded molecular surface. This final part demonstrates how the changes in the dimer interface depicted in the first portion of the movie relate to changes of the rotatory axis between the monomers.

**Movie 7.mp4**

**Movie 7. Catalytic and structural heterogeneity in purified dimeric bovine ATP synthase**. This 38 sec movie shows all the motions observed between monomers accompanying changes in rotational state, and flexions of monomers in fixed rotational states, and combinations of both. The same sequence of reconstructions appear three times at increasing speeds to demonstrate the highly dynamic nature of the assembly. From 00:00 sec to 00:18 sec, in a side view of the dimer, the reconstructions are changed at a rate of *ca*. 3 sec^-1^. In the second part from 00:18 sec to 00:26 sec the rate is *ca*. 9 sec^-1^. In the final segment from 27 sec to 38 sec, the rate is *ca*. 24 sec^-1^ and the complexes are viewed from the matrix. All combinations of rotational state and the extents of the relative motions of the monomers are represented.

