## Supplementary Information for "The mobility of interfaces between monomers in dimeric bovine ATP synthase participates in the ultrastructure of inner mitochondrial membranes"

### Materials and Methods

**Purification of dimeric bovine ATP synthase.** The dimers were purified in the presence of glycodiosgenin, Brij-35 and phospholipids, as described elsewhere (Spikes et al., 2020).

**Image processing and analysis of heterogeneity.** Grids for cryo-em containing the dimeric bovine ATP synthase were prepared and imaged by high-resolution cryo-electron microscopy as described before (Spikes et al., 2020). The particle data were sorted into sub-sets in which the catalytic state of each monomer in the pair was defined (Scheme S1) and then further classified to reveal additional conformational sub-states within them (Scheme S2 and Fig. S2). The pre-processing of micrographs, automated particle picking and initial particle selection by 2D classification have been described elsewhere (Spikes et al., 2020) and are summarized below in Scheme S1. All image processing procedures described here were performed using RELION-3.0 (Scheres, 2012a; Scheres, 2012b; Fernandez-Leiro and Scheres, 2017; Zivanov et al., 2018; Zivanov et al., 2020).

Composite dimer models were constructed from published models of bovine ATP synthase including those of the monomeric enzyme, which were built and refined into focussed locally refined cryo-em reconstructions of higher resolution using COOT (Emsley et al., 2010) and Phenix (Afonine et al., 2018; Liebschner et al., 2019) (see Table S1 and (Spikes et al., 2020)). They were rigid body fitted into reconstructions of the dimeric enzyme produced according to Schemes S1 and S2. The surfaces of cryo-em reconstructions were colored according to the subunit assignments in composite dimer models as in Movies 3, 4, 5 and 6. An octomeric assembly of porcine ATP synthase (Fig. S3) was made from a re-interpreted model of porcine

ATP synthase (EMD-0667). Figures and movies were prepared using UCSF ChimeraX (Goddard et al., 2018).

**Table S1. Published structures employed in this work**

| | PBD | EMDB | Description | Reference |
|---|---|---|---|---|
| **State 1** | 6YY0 | EMD-11001 | Bovine ATP synthase catalytic and rotor domains in rotational state 1 | a |
| **State 2** | 6Z1R | EMD-11039 | Bovine ATP synthase catalytic and rotor domains in rotational state 2 | a |
| **State 3** | 6Z1U | EMD-11040 | Bovine ATP synthase catalytic and rotor domains in rotational state 3 | a |
| **$F_o$** | 6ZBB | EMD-11149 | Bovine ATP synthase monomeric membrane domain | a |
| **State 1 composite** | 6ZPO | EMD-11342 | Complete bovine ATP synthase monomer in rotational state 1 | a |
| **State 2 composite** | 6ZQM | EMD-11368 | Complete bovine ATP synthase monomer in rotational state 2 | a |
| **State 3 composite** | 6ZQN | EMD-11369 | Complete bovine ATP synthase monomer in rotational state 3 | a |
| **Porcine $F_o$** | 6ZMR | EMD-0668 | Porcine ATP synthase monomeric membrane domain, reinterpreted | a, b |
| **Porcine $F_o$ tetramer** | 6ZNA | EMD-0667 | Porcine ATP synthase tetrameric membrane domain, reinterpreted | a, b |

a, Spikes et al, 2020; b, Gu et al, 2020

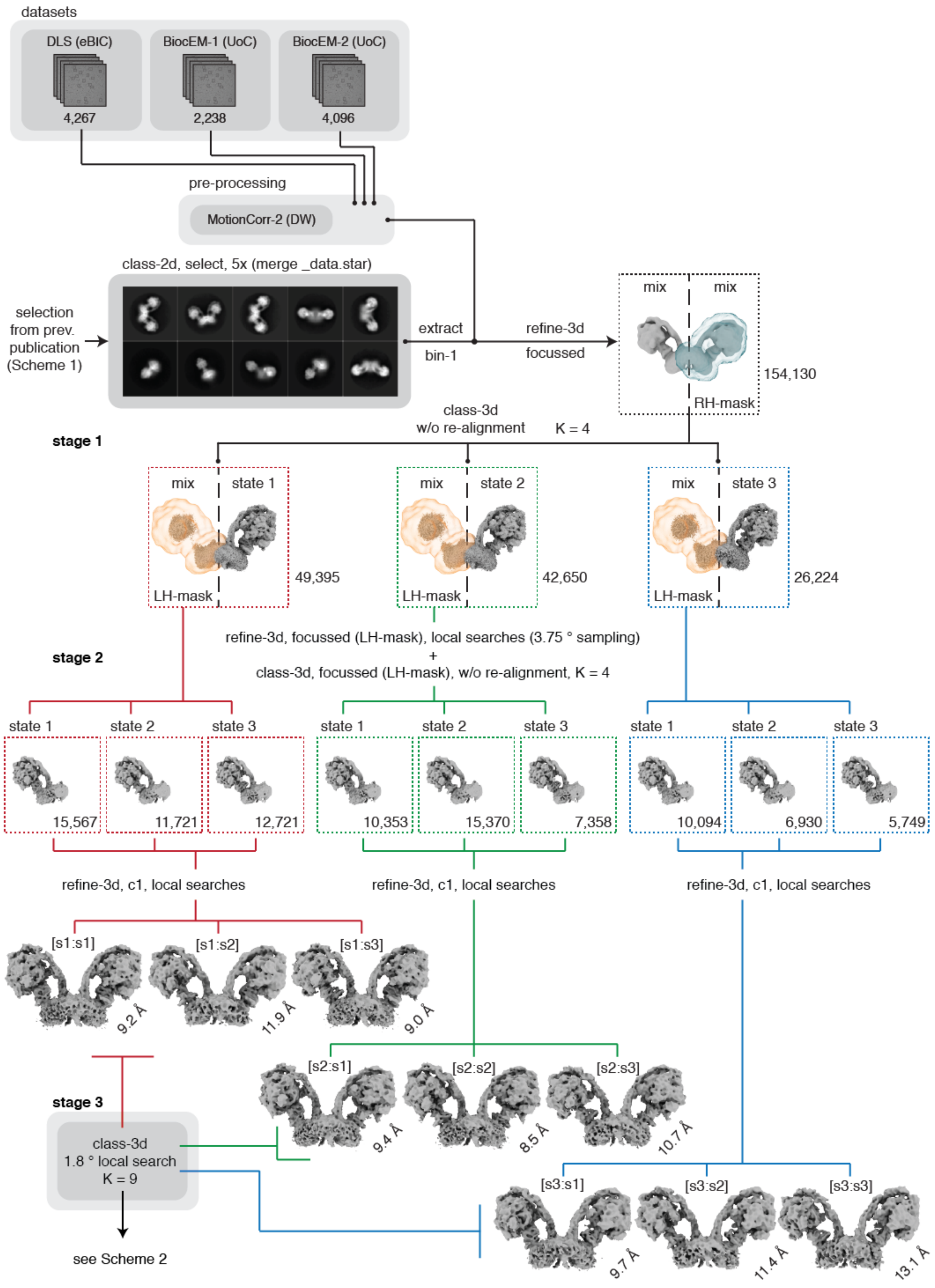


**Scheme S1. Enumeration of all catalytic state combinations in dimeric bovine ATP synthase.** Because the dimer particles in the data-sets were heterogeneous, high resolution structures of the intact dimer could not be determined by refinement of the whole particle set with or without the imposition of c2 symmetry. Therefore, consensus reconstructions represent an average of two distinct types of heterogeneity arising, first from the positions of the asymmetrical $F_1$-domain catalytic relative to the peripheral stalk in the monomeric complex, which are independent of one another, and second from variance in the spatial relationship between monomers arising from differences in the angle between the rotatory axes of the two $F_1$-$c_8$ domains. Therefore, a hierarchical classification strategy employing three stages was applied. First, the catalytic states of each half of the assembly were resolved independently, in order to provide consensus dimer reconstructions in each of the nine combinations of rotational state (shown above), Then they were resolved into classes representing distinct structural conformations with defined catalytic states (Scheme S2). Raw movie frames were corrected for motion, and were dose-weighted to improve the reconstructions in lieu of particle polishing of many individual sub-sets. 154,130 dimer particles, selected by 2D classification in a previous work (Spikes et al., 2020), were re-extracted from the dose-weighted micrographs at a sampling rate of 1.048 Å/pix and separated into rotational states of the right monomer by refining the particles whilst employing a monomeric mask (blue) against the consensus dimer reference, and subsequently classifying without particle re-alignment (stage 1). This procedure yielded 49,395 particles in state-1, 42,650 particles in state-2, 26,224 particles in state 3 and 39,891 particles in a fourth class that were discarded. Each of the three classes representing the rotational states of the right molecule were refined against the consensus dimer reference with a mask encompassing the left monomer (orange). Rotation of the particles about the pseudo-twofold rotational symmetry axis was prevented by local angular searches from the orientations determined for the right monomer, thereby maintaining the relative positions of the right and left monomers with respect to the refinement reference. The left monomer was classified into the three rotational states without particle re-alignment (stage 2) from the refined orientations, leading to particle sub-sets where the rotational state of each monomer in the pair is defined, albeit with a heterogeneous relationship between their spatial arrangement in the overall assembly. The colored dashed boxes contain the number of particles in each sub-state. During this process, an additional 22,826 particles in the fourth class failed to separate into a distinct rotational state and were discarded. As before, rotation of the particles about the pseudo-c2 symmetry axis was prevented by refinement of particle sub-sets without symmetry and local searches from the previously determined orientations. These structures are shown at the bottom of the scheme and are named according to the catalytic state of the right monomer, followed by the left monomer (i.e. the order in which they were classified) as follows; states [s1:s1], [s1:s2], [s1:s3], [s2:s1], [s2:s2], [s2:s3], [s3:s1], [s3:s2] and [s3:s3] at resolutions of 9.2, 11.9, 9.0, 9.4, 8.5, 10.7, 9.7, 11.4, and 13.1 Å, respectively, with applied B-factors ranging from -215 to -500 $Å^2$. In stage 3, each particle set was classified further with reference to its respective consensus structure without particle re-alignment into nine classes (see Scheme S2).

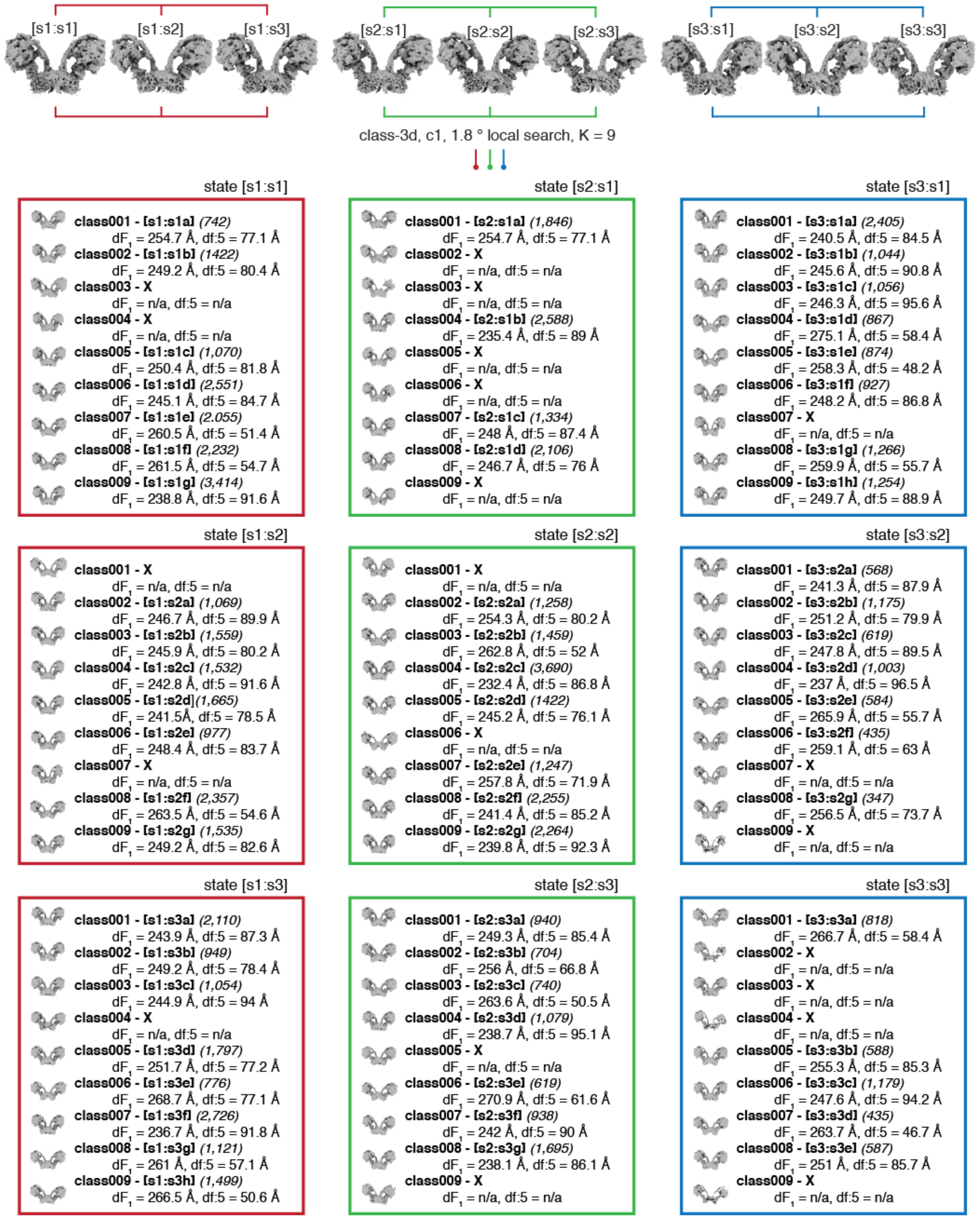


**Scheme S2. Identification and analysis of structural heterogeneity in dimeric bovine ATP synthase.** Each particle set represents the dimers in defined catalytic states (see Scheme S1). Local searches from the consensus orientations for alignment were enforced and the dimer particles in each set were classified into nine additional classes, shown in the colored boxes, right monomers in state 1, state 2 and state 3 in red, green and blue, respectively. The composite atomic models for the corresponding rotational state in each half of the molecule were rigid body fitted into the reconstructions. Two centroids, representing the catalytic domain and residue 5 of subunit f in a small matrix protrusion, were calculated in each monomer and the distances between them in each dimer were measured. This pair of distances, $dF_1$ and df:5, serve as proxy for measurements of the angle between the rotatory axes. These measurements in Ångströms accompany the number of particles (brackets) and a map identifier (square brackets). The per-particle distributions of the measurements are summarised in Fig. S3. Twenty-two maps, with too few particles or with obvious artefacts and denoted with an "X", were excluded from this analysis, leaving fifty-nine reconstructions (see Table S2).

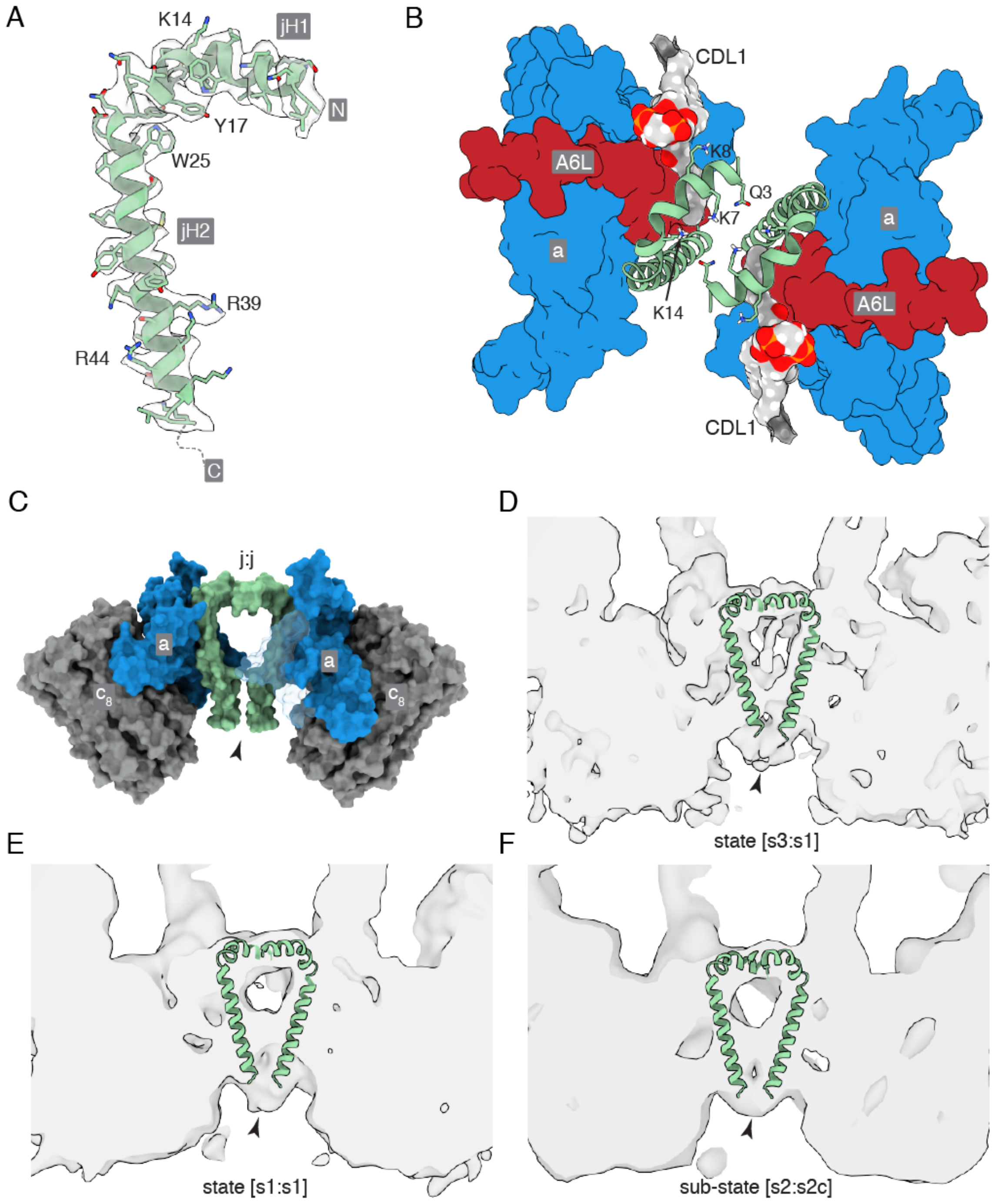


**Fig. S1. The topology of subunit j in dimeric bovine ATP synthase.** *A,* side view of the cryo-em density of the subunit j (grey transparency) and the fitted atomic model (sea-foam green). Residues used to determine the sequence register during model building and their corresponding densities are labelled. The N-terminus of subunit j is folded into an amphipathic α-helix jH1 (residues 1-20), which lies in the lipid head group region on the matrix side of the IMM, followed by the transmembranous jH2 which lies adjacent to A6L and is associated with subunit a. The C-terminal region of jH2 (residues 40-49), protrudes into the IMS, indicated in *C* by the arrowhead; *B*, top view, from inside the mitochondrial matrix between the two peripheral stalks of the dimer (not shown), with the a-subunit (cornflower blue) and A6L-subunit (brick red) shown as a molecular surface and the j-subunit (sea-foam green) in cartoon. The negatively charged headgroup of CDL1 (grey surface, colored by heteroatom) is bound to jK8, and to residues fQ38 and fY42 of the f-subunit in the wedge (not shown) and residues aT33, and A6LK27 and A6LK30. Residues jQ3, jK7 and jK14 are shown projecting toward the interface; *C*, side view, rotated 90° into the plane of the paper with respect to *B*, of the molecular surfaces of the a-subunit (cornflower blue) and the j-subunit (sea-foam green), and the $c_8$-ring (grey), in the dimeric membrane domain of the bovine ATP synthase; *D-F,* cross-sectional side views of three bovine ATP synthase dimers that exemplify density, attributed to the C-terminal regions of the j subunits, beyond the modelled residues. The state [s3:s1], state [s1:s1] and sub-state [s2:s2c] dimers are shown in *D*, *E* and *F,* respectively. The modelled regions of the j-subunits are overlayed onto the transparent density.

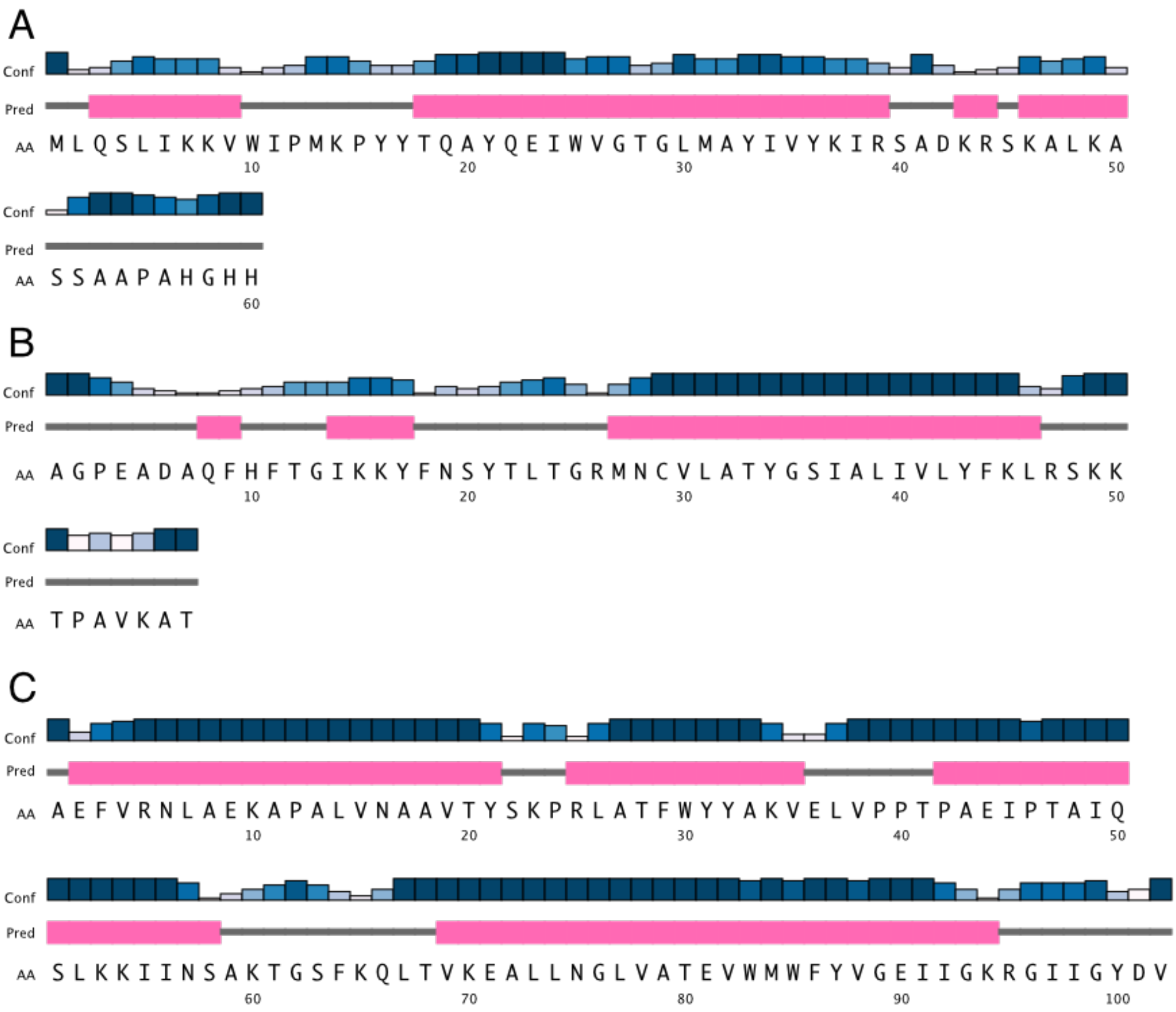


**Fig. S2. The predicted secondary structures of subunits j, k and g of bovine ATP synthase.** In *A*, *B* and *C*, the sequences of subunits j, k and g, respectively, are given in single letter amino acid code, and their predicted secondary structure elements and per-residue confidence scores are shown, respectively, in the lower, middle and upper rows of each panel. α-Helices are pink, and extended structures grey. Confidence scores are shown on a white-blue color scale, with dark blue being the most confident. The secondary structure predictions were performed with PSIPRED (Buchan et al., 2013; Buchan and Jones, 2019). In the structure of bovine subunit j, residues 1-20 form an amphipathic α-helix (jH1) that lies in the plane of the IMM side leaflet and residues j22-39 form the transmembrane region of jH2. The C-terminal region of jH2 (residues 40-49) protrudes into the IMS. The rest of the sequence, comprising residues j50-60, was not modelled but is predicted to form an extended structure as shown above.

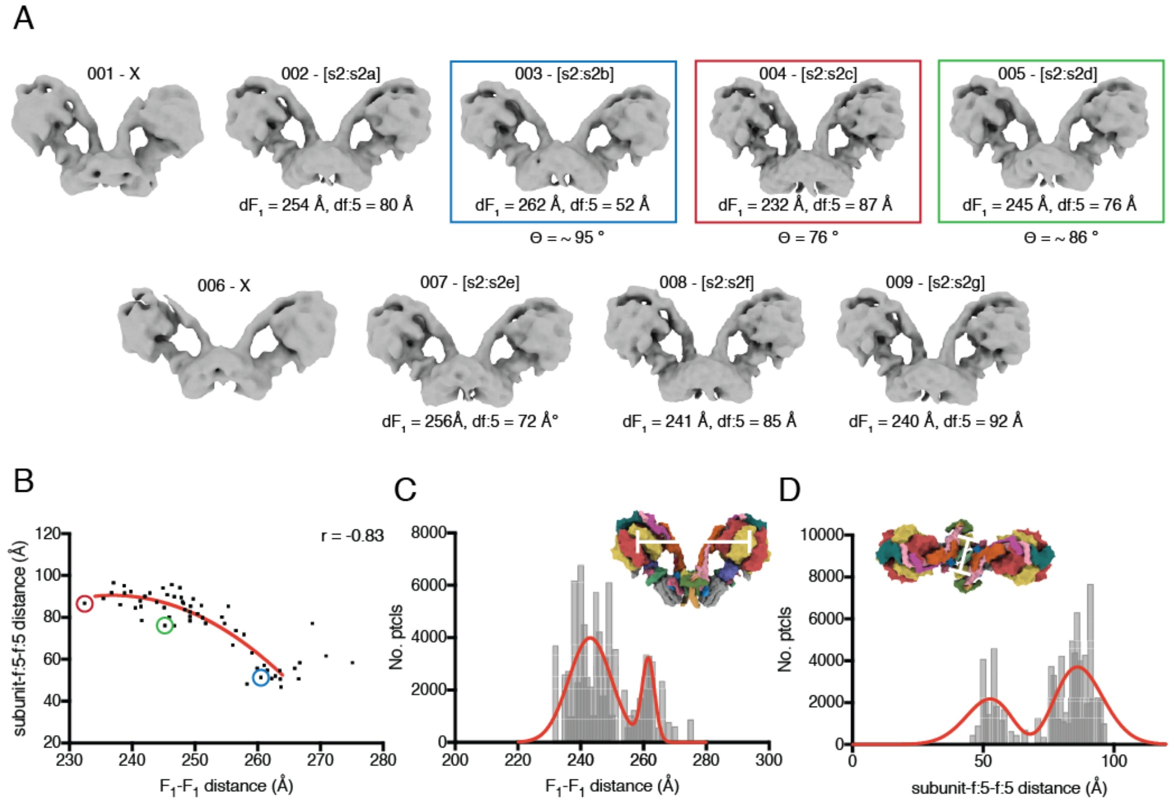


**Fig. S3. Quantitation of the structural heterogeneity observed in dimeric bovine ATP synthase.** *A*, stage 3 of the hierarchical classification (see Schemes 1 and 2) for dimer particles in catalytic state [s2:s2]. Particles in class 001 and 006 were discarded; Movies 3, and 4 and 5, are derived from reconstructions of sub-states [s2:s2c], [s2:s2f] and [s2:s2g], and [s2:s2c], [s2:s2a], [s2:s2e] and [s2:s2b], respectively; *B*, correlation between the distance between $F_1$-domains and the width of the membrane. Encircled dots correspond to the maps in colored boxes in *A*. As the angle between the rotatory axes increases, the catalytic domains separate requiring the dimer interface to re-arrange, leading to the membrane domain becoming narrower in the direction perpendicular to the plane of the rotatory axes (as in *D, Inset*). *C* and *D*, distribution of the distances between $F_1$-domains and between residues 5 in f-subunits, respectively. The most abundant inter-$F_1$ distances clustered at *ca.* 240 Å, and the distance between residues 5 in f-subunits was *ca.* 80-90 Å, which corresponds to a range of angle between rotatory axes of *ca.* 76-86°. In *C* and *D*, the insets show the approximate points of measurement.

**Production of movies**

**Movie 1.** **Structural heterogeneity in dimeric bovine ATP synthase associated with catalysis.** Two copies of the atomic model of the bovine membrane domain (6ZBB) were rigid body fitted into the cryo-em density of the state [s2:s2] consensus dimer reconstruction and a simulated 7 Å density was produced from both chains of subunits a and j. Then, each of the other consensus dimer reconstructions produced by stage 2 of the hierarchical classification procedure (Scheme S1) were fitted to this simulated density so that the maps were aligned by the membrane domains, and the movie was produced by cross-fading between the different reconstructions in the following order; state [s1:s3], [s3:s1], [s1:s2], [s2:s3], [s3:s2], [s2:s1], [s1:s1], [s2:s2], [s3:s3], [s1:s1].

**Movie 2. The rotary cycle during synthesis and hydrolysis.** This movie was produced in the same manner as Movie 1 except that the composite atomic model of the corresponding rotational state, 6ZPO, 6ZQM and 6ZQN (ref) for states 1, 2 and 3, respectively, in each monomer was fitted into the consensus reconstruction and the surface of each subunit was colored accordingly.

**Movie 3. Pivoting of the membrane domains of adjacent monomers of bovine ATP synthase about the matrix contact between j-subunits during catalysis.** This movie was produced in the same manner as Movies 1 and 2 except that the composite atomic model of the corresponding rotational state, 6ZPO, 6ZQM and 6ZQN (ref) for states 1, 2 and 3, respectively, in each monomer was fitted into the appropriate consensus reconstruction and the surfaces of the cryo-em densities were hidden. Side chains were hidden from the atomic model, and the secondary structure elements were dilated in appearance. The movie was produced by cross-fading between the different models in the order state [s1:s2] > [s2:s3] > [s3:s1] > [s1:s2] and as described in the legend of Movie 3.

**Movie 4. The fluidity, independent of catalysis, of the monomer:monomer interface at rotatory axis angles less than or equal to 90°.** Composite atomic models of the bovine ATP synthase monomer in rotational state 2 (6ZQM (ref)) were rigid body fitted into the dimer sub-state [s2:s2c] reconstruction, in which the angle between rotatory axes is *ca.* 76°, produced by stage 3 of the hierarchical classification (see Schemes S1 and S2). A simulated 7 Å density was created from the model of the left monomer (with respect to the screen viewing direction) to which were fitted the reconstructions of the [s2:s2f] and [s2:s2g] dimer sub-states, aligning the reconstructions to a single monomer. Copies of the state 2 atomic model (6ZQM (ref)) were fitted into the remaining monomer domains and the surfaces of individual subunits were colored according to their position in the atomic model.

**Movie 5. Trajectory, independent of catalysis, towards the formation of a wide angle between the central axes of the $F_1$-$c_8$ domains in a dimer of ATP synthase.** Composite atomic models of the bovine ATP synthase monomer in rotational state 2 (6ZQM (ref)) were rigid body fitted into the sub-state [s2:s2c] reconstruction, in which the angle between rotatory axes is *ca.* 76°, produced by stage 3 of the hierarchical classification (see Schemes S1 and S2). A simulated 7 Å density was created from both chains of subunit a and subunit j to which were fitted the reconstructions of the [s2:s2a], [s2:s2e] and [s2:s2b] dimer sub-states, to align them by their membrane domains. Then each reconstruction was re-sampled onto the coordinate system of the [s2:s2c] sub-state and the densities were interpolated in the order sub-state [s2:s2c] > sub-state [s2:s2a] > sub-state [s2:s2e] > sub-state [s2:s2b]. Copies of the state 2 composite atomic model (6ZQM (ref)) were fitted into the new positions of the sub-state reconstructions, and the positions of the atomic coordinates of each model were similarly interpolated over an equal number of frames, in the same order. In each frame of the interpolated trajectory, the surfaces of subunits were colored in the reconstructions.

**Movie 6. Detailed view of the interface rearrangement, independent of catalysis, arising in the trajectory towards the wide angle dimeric ATP synthase.** Top view, as viewed from the matrix, of subunit a and residues 19-49 of subunit j progressing along the interpolated atomic model trajectory from sub-state [s2:s2c] > [s2:s2a] > [s2:s2e] > [s2:s2b]. The movie was prepared as described in the Supplementary Information, Production of Movies, for Movie 5.

**Movie 7. Catalytic and structural heterogeneity in purified dimeric bovine ATP synthase.** All sub-states identified by the hierarchical classification strategy described in Scheme S2 are shown. They were fitted manually to one another by aligning them approximately via their membrane domains. They are cycled in alphanumerical order starting from sub-state [s1:s1a].

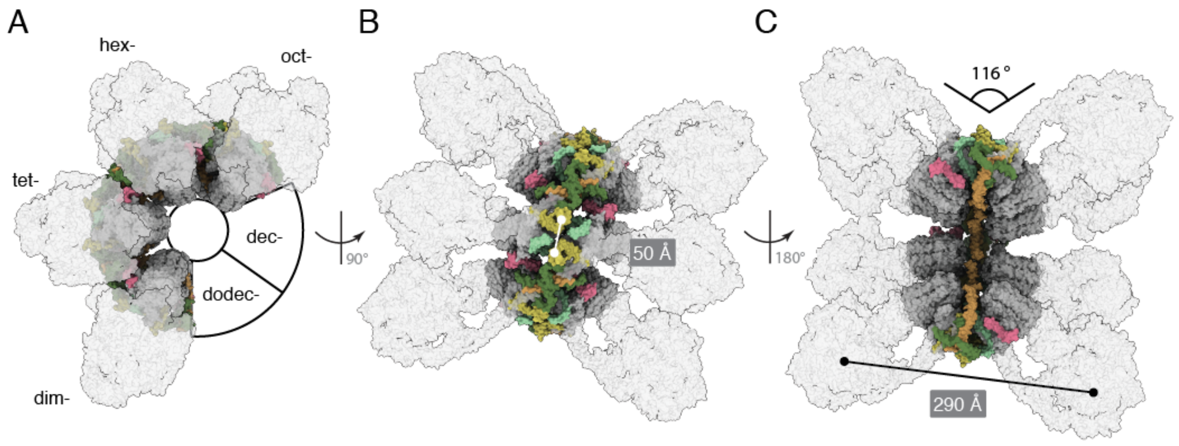


**Fig. S4. Model of a porcine ATP synthase oligomer comprised of the deposited atomic structure of the porcine ATP synthase tetramer.** An octomer of the porcine ATP synthase, reinterpreted as in (Spikes et al., 2020), was constructed by aligning two overlapping tetramers with one another and repeating the process for four dimeric units whilst maintaining the spatial relationship observed in the reported tetrameric assembly in EMBD-0667 (Gu et al., 2019). *A*, side view of the oligomer which would form a closed loop within six dimeric units if the compact tetramer interface is strictly maintained. We suggest that the formation of a tetramer and therefore higher oligomers, in the presence of dimeric inhibitor protein and the absence of a native membrane environment, in this way also contributes to the large $F_1$-$F_1$ distance, narrow membrane domain foot print and very wide rotatory axis observed in the porcine tetramer structure. These things are closely related to the specific membrane domain organization at the interface between monomers. *B*, top view of the oligomer, with the subunit-f:5-subunit-f:5 distance labelled. *C*, a 180° rotated view of *C* with the $F_1$-$F_1$ distance labelled. These measurements can be compared with those in Scheme 2 and Fig. S2 as a proxy for the rotatory axis angle, which was measured as *ca.* 116° by the method described in Fig. 1, but was not measured for all bovine sub-states.

**Table S2. Deposited data-sets relating to the structure of the dimeric bovine ATP synthase**

| | EMDB | Resolution (Å) | B-factor (Å$^2$) | Num. particles | Detail | Comment |
|---|---|---|---|---|---|---|
| **State [s1:s1]** | EMD-11428 | 9.20 | -300 [1] | 15,567 | Hierarchical classification of all dimer particles (Scheme 1) | Consensus, catalytically homogenous, structurally heterogenous |
| **State [s1:s2]** | EMD-11429 | 11.9 | -300 [1] | 11,721 | 〃 | 〃 |
| **State [s1:s3]** | EMD-11430 | 9.00 | -260 [1] | 12,721 | 〃 | 〃 |
| **State [s2:s1]** | EMD-11431 | 9.40 | -255 [1] | 10,353 | 〃 | 〃 |
| **State [s2:s2]** | EMD-11432 | 8.50 | -215 [1] | 15,370 | 〃 | 〃 |
| **State [s2:s3]** | EMD-11433 | 10.7 | -224 [1] | 7,358 | 〃 | 〃 |
| **State [s3:s1]** | EMD-11434 | 9.70 | -350 [1] | 10,094 | 〃 | 〃 |
| **State [s3:s2]** | EMD-11435 | 11.4 | -265 [1] | 6,930 | 〃 | 〃 |
| **State [s3:s3]** | EMD-11436 | 13.1 | -500 [1] | 5,749 | 〃 | 〃 |
| **Sub-state [s1:s1a]** | EMD-11448 | 20.1 | unsharpened [2] | 742 | Classification of state [s1:s1] consensus particles, K = 9, class001 (Scheme 2) | wide rotatory axis angle, subunit j not resolved |

| | | | | | | |
|---|---|---|---|---|---|---|
| **Sub-state [s1:s1b]** | EMD-11449 | 17.5 | 〃 | 1,422 | Classification of state [s1:s1] consensus particles, K = 9, class002 (Scheme 2) | intermediate rotatory axis angle, subunit j partially resolved |
| **Sub-state [s1:s1c]** | EMD-11450 | 18.1 | 〃 | 1,070 | Classification of state [s1:s1] consensus particles, K = 9, class005 (Scheme 2) | narrow rotatory axis angle, subunit j C-terminus resolved |
| **Sub-state [s1:s1d]** | EMD-11451 | 14.9 | 〃 | 2,551 | Classification of state [s1:s1] consensus particles, K = 9, class006 (Scheme 2) | intermediate rotatory axis angle, subunit j C-terminus resolved |
| **Sub-state [s1:s1e]** | EMD-11452 | 16.4 | 〃 | 2,055 | Classification of state [s1:s1] consensus particles, K = 9, class007 (Scheme 2) | wide rotatory axis angle, subunit j not resolved |
| **Sub-state [s1:s1f]** | EMD-11453 | 15.9 | 〃 | 2,232 | Classification of state [s1:s1] consensus particles, K = 9, class008 (Scheme 2) | wide rotatory axis angle, subunit j not resolved |
| **Sub-state [s1:s1g]** | EMD-11454 | 13.8 | 〃 | 3,414 | Classification of state [s1:s1] consensus particles, K = 9, class009 (Scheme 2) | narrow rotatory axis angle, subunit j C-terminus resolved |
| **Sub-state [s1:s2a]** | EMD-11460 | 17.5 | unsharpened | 1,069 | Classification of state [s1:s2] consensus particles, K = 9, class002 (Scheme 2) | narrow rotatory axis angle, subunit j C-terminus resolved |
| **Sub-state [s1:s2b]** | EMD-11461 | 16.9 | 〃 | 1,559 | Classification of state [s1:s2] consensus particles, K = 9, class003 (Scheme 2) | intermediate rotatory axis angle, subunit j C-terminus resolved |

| | | | | | | |
|---|---|---|---|---|---|---|
| **Sub-state [s1:s2c]** | EMD-11462 | 16.9 | 〃 | 1,532 | Classification of state [s1:s2] consensus particles, K = 9, class004 (Scheme 2) | narrow rotatory axis angle, subunit j C-terminus resolved |
| **Sub-state [s1:s2d]** | EMD-11463 | 16.9 | 〃 | 1,665 | Classification of state [s1:s2] consensus particles, K = 9, class005 (Scheme 2) | narrow rotatory axis angle, subunit j C-terminus resolved |
| **Sub-state [s1:s2e]** | EMD-11464 | 18.7 | 〃 | 977 | Classification of state [s1:s2] consensus particles, K = 9, class006 (Scheme 2) | intermediate rotatory axis angle, subunit j C-terminus resolved, partial occupancy |
| **Sub-state [s1:s2f]** | EMD-11465 | 15.9 | 〃 | 2,357 | Classification of state [s1:s2] consensus particles, K = 9, class008 (Scheme 2) | wide rotatory axis angle, subunit j not resolved |
| **Sub-state [s1:s2g]** | EMD-11466 | 16.9 | 〃 | 1,535 | Classification of state [s1:s2] consensus particles, K = 9, class009 (Scheme 2) | intermediate rotatory axis angle, subunit j C-terminus resolved |
| **Sub-state [s1:s3a]** | EMD-11472 | 15.9 | unsharpened | 2,110 | Classification of state [s1:s3] consensus particles, K = 9, class001 (Scheme 2) | intermediate rotatory axis angle, subunit j C-terminus resolved |
| **Sub-state [s1:s3b]** | EMD-11473 | 18.7 | 〃 | 949 | Classification of state [s1:s3] consensus particles, K = 9, class002 (Scheme 2) | intermediate rotatory axis angle, subunit j C-terminus resolved not resolved |
| **Sub-state [s1:s3c]** | EMD-11474 | 18.7 | 〃 | 1,054 | Classification of state [s1:s3] consensus particles, K = 9, class003 (Scheme 2) | intermediate rotatory axis angle, subunit j C-terminus resolved, partial occupancy |

| | | | | | | |
|---|---|---|---|---|---|---|
| **Sub-state [s1:s3d]** | EMD-11475 | 15.9 | 〃 | 1,797 | Classification of state [s1:s3] consensus particles, K = 9, class005 (Scheme 2) | intermediate rotatory axis angle, subunit j C-terminus resolved |
| **Sub-state [s1:s3e]** | EMD-11476 | 20.2 | 〃 | 776 | Classification of state [s1:s3] consensus particles, K = 9, class006 (Scheme 2) | intermediate rotatory axis angle, subunit j C-terminus resolved not resolved |
| **Sub-state [s1:s3f]** | EMD-11477 | 15.4 | 〃 | 2,726 | Classification of state [s1:s3] consensus particles, K = 9, class007 (Scheme 2) | narrow rotatory axis angle, subunit j C-terminus resolved |
| **Sub-state [s1:s3g]** | EMD-11479 | 18.1 | 〃 | 1,121 | Classification of state [s1:s3] consensus particles, K = 9, class008 (Scheme 2) | wide rotatory axis angle, subunit j not resolved |
| **Sub-state [s1:s3h]** | EMD-11480 | 16.9 | 〃 | 1,499 | Classification of state [s1:s3] consensus particles, K = 9, class009 (Scheme 2) | wide rotatory axis angle, subunit j not resolved |
| **Sub-state [s2:s1a]** | EMD-11484 | 16.4 | unsharpened | 1,846 | Classification of state [s2:s1] consensus particles, K = 9, class001 (Scheme 2) | wide rotatory axis angle, subunit j not resolved |
| **Sub-state [s2:s1b]** | EMD-11485 | 15.4 | 〃 | 2,588 | Classification of state [s2:s1] consensus particles, K = 9, class004 (Scheme 2) | narrow rotatory axis angle, subunit j C-terminus resolved |
| **Sub-state [s2:s1c]** | EMD-11486 | 17.5 | 〃 | 1,334 | Classification of state [s2:s1] consensus particles, K = 9, class007 (Scheme 2) | intermediate rotatory axis angle, subunit j C-terminus resolved |

| | | | | | | |
|---|---|---|---|---|---|---|
| **Sub-state [s2:s1d]** | EMD-11487 | 14.9 | 〃 | 2,106 | Classification of state [s2:s1] consensus particles, K = 9, class008 (Scheme 2) | narrow rotatory axis angle, subunit j C-terminus resolved |
| **Sub-state [s2:s2a]** | EMD-11499 | 17.5 | unsharpened | 1,258 | Classification of state [s2:s2] consensus particles, K = 9, class002 (Scheme 2) | wide rotatory axis angle, subunit j partially resolved |
| **Sub-state [s2:s2b]** | EMD-11500 | 17.5 | 〃 | 1,459 | Classification of state [s2:s2] consensus particles, K = 9, class003 (Scheme 2) | wide rotatory axis angle, subunit j not resolved |
| **Sub-state [s2:s2c]** | EMD-11501 | 12.8 | 〃 | 3,690 | Classification of state [s2:s2] consensus particles, K = 9, class004 (Scheme 2) | narrow rotatory axis angle, subunit j C-terminus resolved, most homogenous |
| **Sub-state [s2:s2d]** | EMD-11502 | 16.4 | 〃 | 1,422 | Classification of state [s2:s2] consensus particles, K = 9, class005 (Scheme 2) | intermediate rotatory axis angle, subunit j C-terminus resolved, partial occupancy |
| **Sub-state [s2:s2e]** | EMD-11503 | 18.1 | 〃 | 1,247 | Classification of state [s2:s2] consensus particles, K = 9, class007 (Scheme 2) | wide rotatory axis angle, subunit j not resolved |
| **Sub-state [s2:s2f]** | EMD-11504 | 14.9 | 〃 | 2,255 | Classification of state [s2:s2] consensus particles, K = 9, class008 (Scheme 2) | intermediate rotatory axis angle, subunit j C-terminus resolved |
| **Sub-state [s2:s2g]** | EMD-11505 | 14.9 | 〃 | 2,264 | Classification of state [s2:s2] consensus particles, K = 9, class009 (Scheme 2) | intermediate rotatory axis angle, subunit j C-terminus resolved |

| | | | | | | |
|---|---|---|---|---|---|---|
| **Sub-state [s2:s3a]** | EMD-11506 | 18.1 | unsharpened | 940 | Classification of state [s2:s3] consensus particles, K = 9, class001 (Scheme 2) | intermediate rotatory axis angle, subunit j C-terminus resolved |
| **Sub-state [s2:s3b]** | EMD-11507 | 19.4 | 〃 | 704 | Classification of state [s2:s3] consensus particles, K = 9, class002 (Scheme 2) | intermediate rotatory axis angle, subunit j C-terminus not resolved |
| **Sub-state [s2:s3c]** | EMD-11508 | 20.2 | 〃 | 740 | Classification of state [s2:s3] consensus particles, K = 9, class003 (Scheme 2) | wide rotatory axis angle, subunit j not resolved |
| **Sub-state [s2:s3d]** | EMD-11509 | 18.1 | 〃 | 1,079 | Classification of state [s2:s3] consensus particles, K = 9, class004 (Scheme 2) | narrow rotatory axis angle, subunit j C-terminus resolved |
| **Sub-state [s2:s3e]** | EMD-11510 | 20.9 | 〃 | 619 | Classification of state [s2:s3] consensus particles, K = 9, class006 (Scheme 2) | wide rotatory axis angle, subunit j not resolved |
| **Sub-state [s2:s3f]** | EMD-11511 | 18.7 | 〃 | 938 | Classification of state [s2:s3] consensus particles, K = 9, class007 (Scheme 2) | narrow rotatory axis angle, subunit j C-terminus partially resolved |
| **Sub-state [s2:s3g]** | EMD-11512 | 16.4 | 〃 | 1,695 | Classification of state [s2:s3] consensus particles, K = 9, class008 (Scheme 2) | narrow rotatory axis angle, subunit j C-terminus resolved |
| **Sub-state [s3:s1a]** | EMD-11527 | 15.4 | unsharpened | 2,405 | Classification of state [s3:s1] consensus particles, K = 9, class001 (Scheme 2) | narrow rotatory axis angle, subunit j C-terminus resolved |

| | | | | | | |
|---|---|---|---|---|---|---|
| **Sub-state [s3:s1b]** | EMD-11528 | 18.1 | 〃 | 1,044 | Classification of state [s3:s1] consensus particles, K = 9, class002 (Scheme 2) | intermediate rotatory axis angle, subunit j C-terminus resolved |
| **Sub-state [s3:s1c]** | EMD-11529 | 18.1 | 〃 | 1,056 | Classification of state [s3:s1] consensus particles, K = 9, class003 (Scheme 2) | intermediate rotatory axis angle, subunit j C-terminus partially resolved |
| **Sub-state [s3:s1d]** | EMD-11530 | 18.7 | 〃 | 867 | Classification of state [s3:s1] consensus particles, K = 9, class004 (Scheme 2) | wide rotatory axis angle, subunit j not resolved |
| **Sub-state [s3:s1e]** | EMD-11531 | 19.4 | 〃 | 874 | Classification of state [s3:s1] consensus particles, K = 9, class005 (Scheme 2) | wide rotatory axis angle, subunit j not resolved |
| **Sub-state [s3:s1f]** | EMD-11532 | 18.7 | 〃 | 927 | Classification of state [s3:s1] consensus particles, K = 9, class006 (Scheme 2) | intermediate rotatory axis angle, subunit j C-terminus partially resolved |
| **Sub-state [s3:s1g]** | EMD-11533 | 17.5 | 〃 | 1,266 | Classification of state [s3:s1] consensus particles, K = 9, class008 (Scheme 2) | wide rotatory axis angle, subunit j not resolved |
| **Sub-state [s3:s1h]** | EMD-11534 | 17.5 | 〃 | 1,254 | Classification of state [s3:s1] consensus particles, K = 9, class009 (Scheme 2) | narrow rotatory axis angle, subunit j C-terminus partially resolved |
| **Sub-state [s3:s2a]** | EMD-11535 | 16.9 | unsharpened | 568 | Classification of state [s3:s2] consensus particles, K = 9, class001 (Scheme 2) | narrow rotatory axis angle, subunit j C-terminus resolved |

| | | | | | | |
|---|---|---|---|---|---|---|
| **Sub-state [s3:s2b]** | EMD-11536 | 20.2 | 〃 | 1,175 | Classification of state [s3:s2] consensus particles, K = 9, class002 (Scheme 2) | intermediate rotatory axis angle, subunit j C-terminus partially resolved |
| **Sub-state [s3:s2c]** | EMD-11537 | 18.1 | 〃 | 619 | Classification of state [s3:s2] consensus particles, K = 9, class003 (Scheme 2) | intermediate rotatory axis angle, subunit j C-terminus resolved |
| **Sub-state [s3:s2d]** | EMD-11538 | 20.9 | 〃 | 1,003 | Classification of state [s3:s2] consensus particles, K = 9, class004 (Scheme 2) | narrow rotatory axis angle, subunit j C-terminus partially resolved |
| **Sub-state [s3:s2e]** | EMD-11539 | 18.7 | 〃 | 584 | Classification of state [s3:s2] consensus particles, K = 9, class005 (Scheme 2) | wide rotatory axis angle, subunit j not resolved |
| **Sub-state [s3:s2f]** | EMD-11540 | 20.9 | 〃 | 435 | Classification of state [s3:s2] consensus particles, K = 9, class006 (Scheme 2) | wide rotatory axis angle, subunit j not resolved |
| **Sub-state [s3:s2g]** | EMD-11541 | 20.9 | 〃 | 347 | Classification of state [s3:s2] consensus particles, K = 9, class008 (Scheme 2) | intermediate rotatory axis angle, subunit j C-terminus partially resolved |
| **Sub-state [s3:s3a]** | EMD-11542 | 19.4 | unsharpened | 818 | Classification of state [s3:s3] consensus particles, K = 9, class001 (Scheme 2) | wide rotatory axis angle, subunit j not resolved |
| **Sub-state [s3:s3b]** | EMD-11543 | 20.9 | 〃 | 588 | Classification of state [s3:s3] consensus particles, K = 9, class005 (Scheme 2) | intermediate rotatory axis angle, subunit j C-terminus not resolved |

| | | | | | | |
|---|---|---|---|---|---|---|
| **Sub-state [s3:s3c]** | EMD-11544 | 17.5 | 〃 | 1,179 | Classification of state [s3:s3] consensus particles, K = 9, class006 (Scheme 2) | narrow rotatory axis angle, subunit j C-terminus resolved |
| **Sub-state [s3:s3d]** | EMD-11545 | 23.8 | 〃 | 435 | Classification of state [s3:s3] consensus particles, K = 9, class007 (Scheme 2) | wide rotatory axis angle, subunit j not resolved |
| **Sub-state [s3:s3e]** | EMD-11546 | 21.8 | 〃 | 587 | Classification of state [s3:s3] consensus particles, K = 9, class008 (Scheme 2) | intermediate rotatory axis angle, subunit j C-terminus not resolved |

[1] The maps were sharpened with an *ad hoc* user defined B-factor in lieu of calculating a value from the half-map data.

[2] Without map sharpening.

**Table S3. Summary of the structural models of subunits g, j, and k**

| Subunit | No. of Residues | Residues modelled |
|---|---|---|
| g | 102 | 20-98 |
| j | 60 | 2-49 |
| k | 57 | 12-47 |
